# TWIST1 drives endothelial-to-mesenchymal-transition to stabilize atherosclerotic plaques

**DOI:** 10.1101/2025.05.19.654847

**Authors:** Blanca Tardajos-Ayllon, Mannekomba Diagbouga, Andreas Edsfeldt, Joanna Kalucka, Jovana Serbanovic-Canic, Emily Chambers, Jiangming Sun, Chrysostomi Gialeli, Mark Dunning, Xiuying Li, Suowen Xu, Sheila E. Francis, M Akiko Mammoto, Michael Simons, Helle F Jørgensen, Isabel Goncalves, Paul C. Evans

## Abstract

Atherosclerosis progresses from fatty streaks to complex plaques, with rupture leading to life-threatening complications. Endothelial-to-mesenchymal transition (EndMT) is associated with advanced atherosclerotic plaques, but its role in plaque progression remains unclear. To investigate this, we analyzed the role of TWIST1, a key EndMT-driving transcription factor, in plaque development. Using single-cell RNA sequencing of atherosclerotic plaques from hypercholesterolemic mice with inducible deletion of *Twist1* from endothelial cells (*Twist1^ECKO^ ApoE^-/-^*) we demonstrate that *Twist1* regulates endothelial cell heterogeneity by promoting cell states expressing EndMT markers. Cell tracking confirmed that *Twist1* induces EndMT in advanced plaques. Mechanistically, we found that TWIST1 contributes to EndMT by promoting endothelial migration and proliferation through the transcriptional coactivator PELP1. Additionally, TWIST1 induces AEBP1 transcription, which upregulates COL4A1 to drive endothelial proliferation. Analyses of murine brachiocephalic plaques show that endothelial *Twist1* promotes plaque growth, collagen deposition, and ACTA2-positive cell accumulation, hallmarks of plaque stability, while reducing necrosis and macrophage infiltration, features of plaque instability. Moreover, TWIST1 expression was associated with asymptomatic human carotid atherosclerosis and predicts favourable clinical outcomes. These findings challenge the prevailing view that EndMT destabilizes plaques, by suggesting that TWIST1-driven EndMT can promote plaque stability, offering new insights into atherosclerosis pathophysiology and therapeutic potential.

## INTRODUCTION

Atherosclerosis, a leading cause of mortality worldwide, is a progressive arterial disease that begins as fatty streaks and advances to complex plaques capable of rupture^1^. Disturbed flow (DF) of blood accelerates the transition of plaques into rupture-prone forms characterized by large necrotic cores, macrophage accumulation, reduced vascular smooth muscle cells, and diminished collagen content^2^. This transformation compromises plaque integrity, elevating the risk of rupture-associated complications such as stroke, unstable angina, and myocardial infarction.

Arterial endothelial cells (ECs), typically organized as a monolayer at the luminal surface, play a critical role in maintaining vascular homeostasis by regulating the influx of inflammatory cells and lipoproteins. However, ECs exhibit phenotypic plasticity, allowing them to adapt their functions and characteristics. One such transformation is endothelial-to-mesenchymal transition (EndMT), during which ECs lose markers like CDH5 (VE-cadherin) while acquiring mesenchymal markers such as ACTA2 (alpha-smooth muscle actin)^3–5^. EndMT endows ECs with migratory and invasive properties, enabling their movement from the lumen into underlying tissues where they produce collagen and display myofibroblast-like features ^4,5^. This process plays a fundamental role in cardiovascular development, including the formation of valve leaflets from subendothelial cushions during embryogenesis^6,7^.

EndMT has emerged as a key feature of atherosclerosis, where it is driven by proatherogenic stimuli including DF^8,9^ and pro-inflammatory cytokines^10,11^. Single-cell RNA sequencing (scRNA-seq) coupled with cell-tracking technologies have provided compelling evidence of EndMT and other forms of EC plasticity, such as endothelial-to-inflammatory cell transformation (EndIC)^12^, in advanced atherosclerotic plaques in mice^10,11,13–17^. Several studies have demonstrated a tight correlation between EndMT and atherosclerosis progression; EC-specific deletion of *Frs2a*^10^ and tenascin-X^15^ exacerbate EndMT and worsen atherosclerosis, whereas EC-specific deletion of *Shc* ^16^, *Epsin1*^17^, or *IL-1*^11^ have the opposite effects. However, the precise role of EndMT in atherosclerosis remains incompletely understood.

EndMT is regulated by multiple transcription factors, including TWIST1, which drives EndMT in contexts such as angiogenesis^18^, vascular remodeling^19^, pulmonary hypertension^20^ and fibrosis^21^. Previously, we demonstrated that TWIST1 promotes lipid accumulation in early atherosclerosis ^9^. In this study, we analyse the effects of EC deletion of *Twist1* in pre-existing plaques to further elucidate the role of EndMT in atherosclerosis progression. Our findings reveal that TWIST1 drives EndMT and promotes multiple features of plaque stability. These results challenge the prevailing view that EndMT universally destabilizes plaques, suggesting instead that TWIST1 drives a form of EndMT which may confer beneficial effects in advanced atherosclerosis.

## RESULTS

### *Twist1* regulates EC clusters enriched for EndMT markers

To investigate the role of *Twist1* in regulating endothelial phenotypic changes in advanced atherosclerotic plaques, we performed an EC-specific genetic deletion of *Twist1* in mice with pre-existing plaques (Fig. S1a). This was achieved by feeding *Twist1^ECKO^ ApoE^-/-^* and control mice a Western diet for 14 weeks. Within this period, after 8 weeks, *Twist1* EC deletion was induced by treatment with tamoxifen. Quantitative RT-PCR confirmed tamoxifen-induced deletion of *Twist1* in EC of *Twist1^ECKO^ ApoE^-/-^* mice (Fig. S1b). To examine the impact of Twist1 deletion on plaque EC phenotypes, scRNA-seq was performed on aortas from control and *Twist1^ECKO^ ApoE^-/-^* mice. ECs (CD31+ CD45-) were isolated via enzymatic digestion and FACS sorting before scRNA-seq analysis (Fig. S2a). Following quality control filtering, 1,173 control and 966 Twist1^ECKO^ *ApoE^-/-^* cells were selected for downstream analysis (Fig. S2). Hierarchical clustering analysis identified 11 cell clusters (Fig. 1a). Comparison of *Twist1^ECKO^ ApoE^-/-^* mice and control mice revealed that *Twist1* significantly regulates EC heterogeneity in atherosclerotic plaques (Fig. 1b; Fig. S3). Clusters 2, 7, 8 and 9 were enriched for cells from control mice, while cluster 4 and 5 had a higher proportion of cells from *Twist1^ECKO^ ApoE^-/-^*mice (Fig. 1c). Contributions to clusters 0, 1, 3, 6, and 10 were similar between both groups. These findings suggest that *Twist1* promotes clusters 2, 7, 8 and 9 while suppressing clusters 4 and 5. Given that TWIST1 is induced by DF and suppressed by uniform flow (UF) ^9^, we compared the distribution of DF and UF markers among the clusters. Most of the clusters that were promoted by *Twist1* (clusters 2, 7, 8, 9) exhibited an enrichment of DF markers and a corresponding reduction in UF markers, whereas clusters that were suppressed by *Twist1* (cluster 4 and 5) showed the opposite pattern (Fig. S4). These findings confirm that *Twist1* is a key regulator of EC heterogeneity and responses to DF.

**Figure 1.**
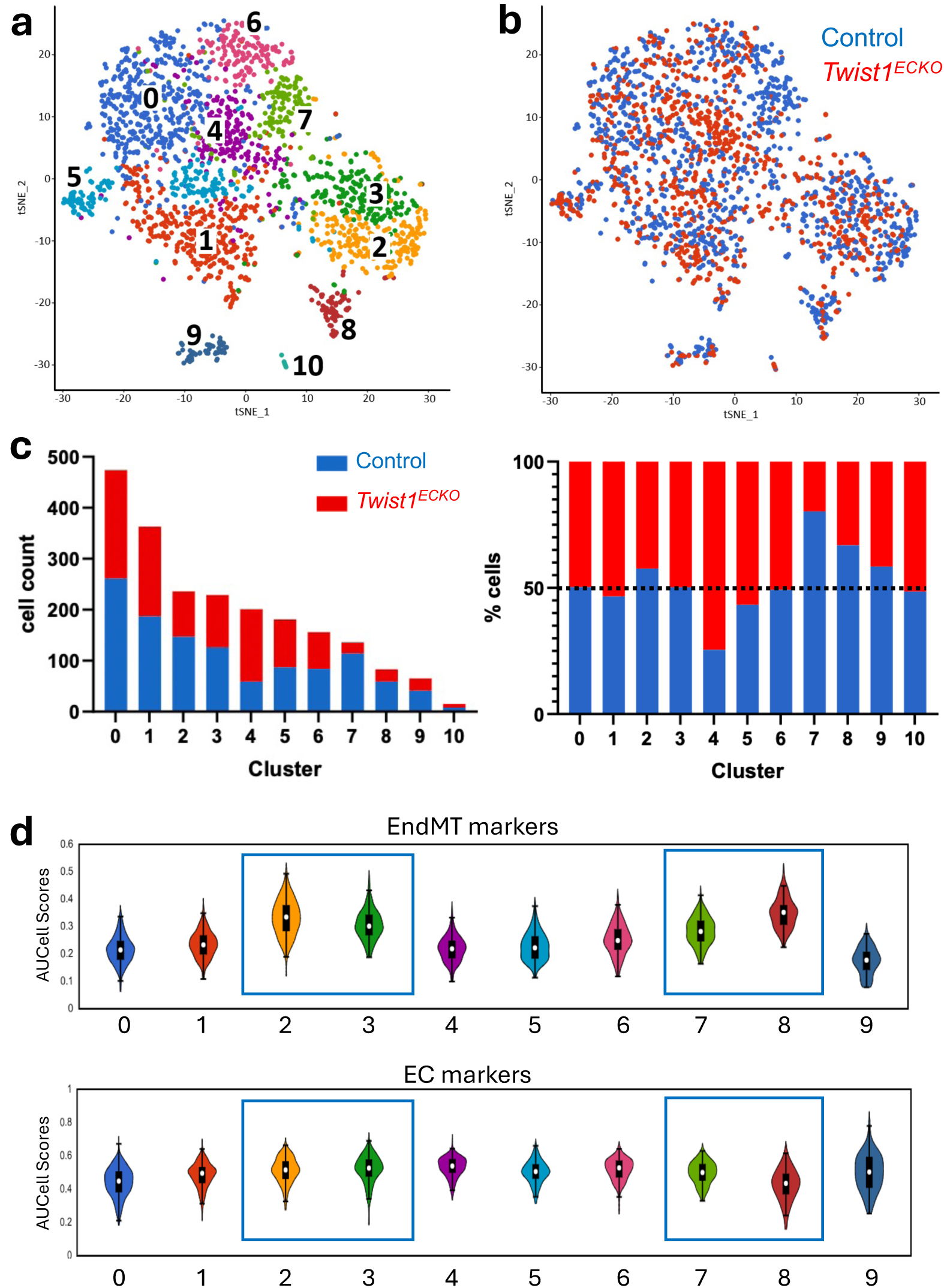
scRNA-seq analysis of *Twist1*-regulated EC heterogeneity. Aortas from *Twist1^ECKO^* and control mice after 14 weeks of Western diet were analysed by FACS of CD31^+^ CD45^−^ cells coupled to scRNA-seq. (A) t-SNE map of single-cell RNA sequencing from *Twist1* EC^KO^ and control mice coloured by cluster assignment. Clusters were identified using unbiased hierarchical clustering. (B) t-SNE map showing the cell contribution of *Twist1* EC^KO^ and control mice to each subpopulation. (C) Bar graphs showing the absolute number of cells from *Twist1* EC^KO^ and control mice in each cluster (left), and the proportion of *Twist1* EC^KO^ and control cells in each cluster (right). The dotted line indicates the 50% threshold. Clusters 2, 7, 8 and 9 are largely composed of ECs derived from control mice, whereas cluster 4 and 5 are mainly composed of ECs derived from *Twist1* EC^KO^ mice. (D) EndMT markers and EC markers were measured in each individual cell in the scRNA-seq dataset and are presented as violin plots as an average (AUCell score). Clusters 2, 3, 7 and 8, which are highlighted with a blue box, show enriched expression of endMT markers.

We conducted an unbiased analysis using cluster-specific nested functional enrichment of gene ontology (GO) terms (Fig. S5). Cluster 10, which contained the fewest cells, was enriched for pathways related to vascular smooth muscle cell (VSMC) signalling and muscle contractility (Fig. S6a). As it likely represented contaminating VSMCs, it was excluded from further analysis. *Twist1*-regulated clusters 2, 4, 5, 7, 8 and 9 were associated with diverse EC functions. Of note, the low frequency cluster 9 is highly enriched for regulators of lipid metabolism and angiogenesis markers and has been previously described^22,23^ (Fig. S6b). It was also noticeable that GO terms related to angiogenesis were observed at higher frequency in *Twist1*-regulated clusters (67%) compared to clusters that were unregulated (0%). Given the close relationship between angiogenesis and EndMT, we next analyzed the distribution of EndMT markers and classical EC markers among clusters (Fig. 1d and Fig. S7). EndMT markers were enriched in clusters 2, 7 and 8 which are *Twist1*-regulated, and cluster 3 which is insensitive to *Twist1*. Cluster 8 showed a striking enrichment of EndMT markers alongside a reduction in EC markers, suggesting that this cluster represents an advanced stage of EndMT. In contrast, clusters 2, 3 and 7 exhibited modest alterations in EC markers suggesting a partial EndMT state. These findings suggest that *Twist1* promotes clusters associated with both advanced (cluster 8) and partial (clusters 2, 7) forms of EndMT in advanced atherosclerotic lesions.

### *Twist1* controls EndMT in advanced atherosclerosis

To directly analyse EndMT, we tracked EC-derived cells in *Cdh5^CreERT^*^2^ *Rosa26^tdTomato^ ApoE^-/-^* mice that were fed a Western diet for 14 weeks to generate lesions. Tamoxifen administration at the 8-week time-point induced EC expression of TdTomato for tracking (Fig. 2a). Tracked EC in the lumen of healthy portions of the brachiocephalic artery expressed the endothelial marker vWF but lacked the EndMT marker ACTA2, indicating a differentiated EC phenotype (Fig. 2b; healthy region labelled H). By contrast, tracked EC within the plaque expressed the EndMT marker ACTA2 but lacked expression of the EC marker vWF, suggesting that they had undergone EndMT (Fig. 2b; plaque region labelled P). *Twist1^ECKO^ ApoE^-/-^* mice exhibited a significant reduction in the accumulation of tracked EC-derived cells within the plaque (Fig. 2c), identifying *Twist1* as a driver of EndMT in atherosclerosis.

**Figure 2.**
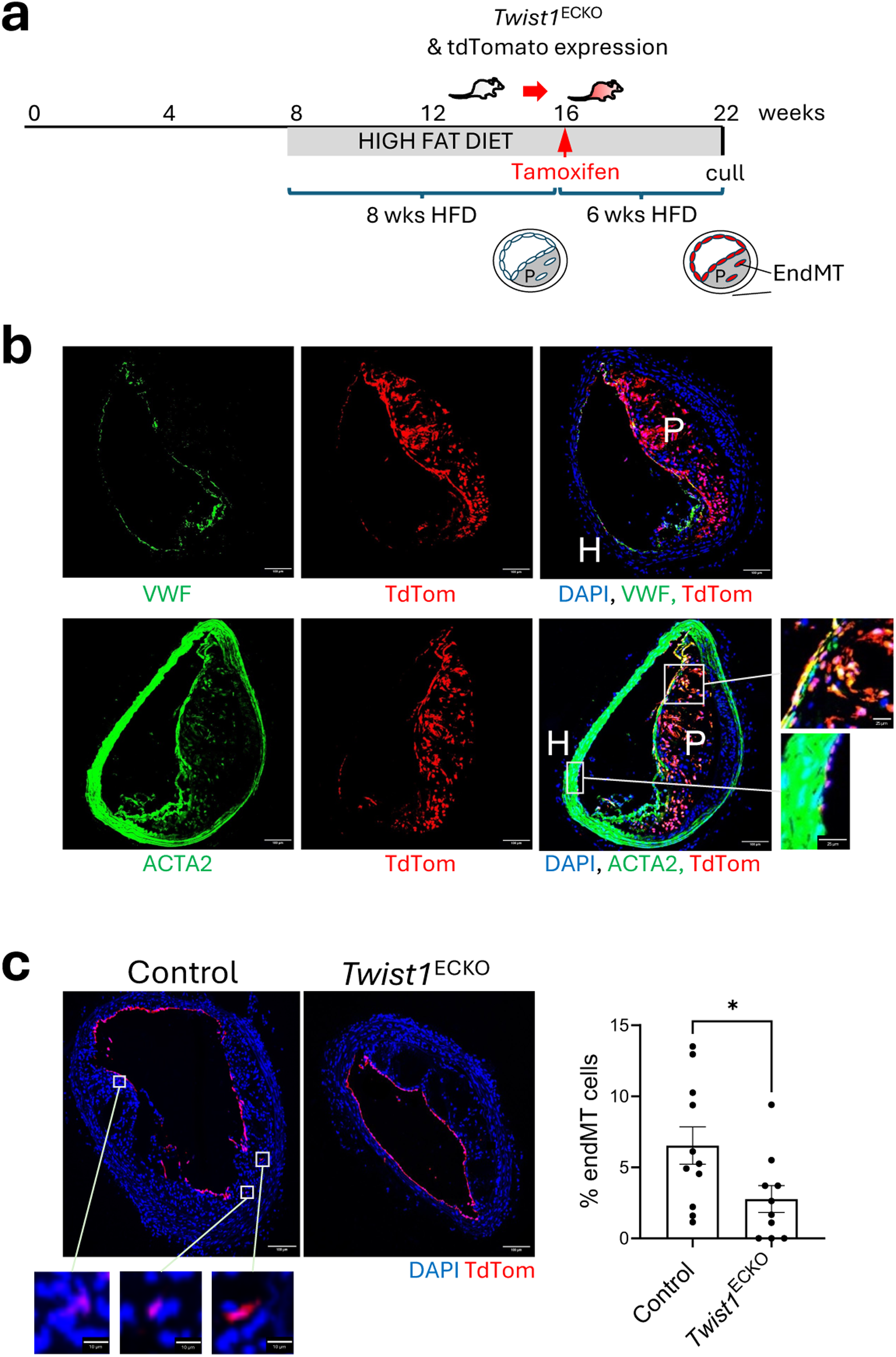
*Twist1* promotes EndMT *in vivo*. (A) Timeline of *Twist1* deletion on an EC-fate mapped background in a model of atherosclerotic progression. Male *Twist1*^ECKO^ (*Twist1^fl/fl^ Cdh5^CreERT^*^2^*^/+^ ApoE^-/-^ Rosa26^TdTomato/TdTomato^*) and control mice (*Twist1^fl/fl^ Cdh5^+/+^ ApoE^-/-^ Rosa26^TdTomato/TdTomato^*) aged 8 weeks were fed a Western diet for 8 weeks to induce atherosclerotic lesions. Tamoxifen was then administered for 5 consecutive days to induce *Twist1* deletion and TdTomato expression in ECs and a Western diet was provided for an additional 6 weeks (totalling 14 weeks of Western diet). (B, C) Brachiocephalic artery atherosclerotic plaques after 14 weeks of Western diet. (B) Frozen sections of brachiocephalic arteries were stained using antibodies against VWF and ACTA2 (green). Rosa26TdTomato^+^ cells are shown in red, and nuclei are counterstained with DAPI (blue). The magnified view of the boxed region shows Rosa26TdTomato/ACTA2 double-positive cells in the plaque and Rosa26TdTomato-positive/ACTA2-negative cells in the luminal area. Representative images are shown (Scale bar=100 μm or 25 μm for magnified views). (C) The percentage of Rosa26TdTomato^+^ cells that have undergone endMT was quantified in *Twist1*^ECKO^ (n=10) and control (n=11) mice. Rosa26TdTomato^+^ cells are shown in red, and nuclei are counterstained with DAPI (blue). The magnified view of the boxed region shows Rosa26TdTomato^+^ cells in the plaque. Representative images are shown (Scale bar=100 μm or 10 μm for magnified views). Mean values are shown +/-standard errors. Differences between means were analysed using an unpaired t-test.

### Molecular mechanisms of TWIST1-driven EndMT

To elucidate the molecular mechanisms underlying TWIST1-driven EndMT, we utilized cultured human aortic EC (HAEC) exposed to DF, conditions known to activate TWIST1 both *in vitro* ^9^ and *in vivo* (Fig. S4). *TWIST1* silencing via siRNA was confirmed (Fig. 3a), and bulk RNA-seq identified 293 genes significantly modulated upon silencing (Fig. 3b). Functional annotation revealed enrichment in gene ontology terms associated with extracellular matrix, development, proliferation, epithelial-to-mesenchymal transition (EMT), and morphogenesis, consistent with a role for TWIST1 in EndMT (Fig. 3c). More specifically, *TWIST1* silencing significantly downregulated EndMT markers while preserving classical endothelial markers (Fig. 3d), suggesting a role in partial EndMT. Functionally, *TWIST1* silencing impaired cell migration (Fig. 3e-3g) and proliferation (Fig. 3h) in HAEC. These findings demonstrate that TWIST1 orchestrates EndMT in HAEC exposed to DF by promoting migration and proliferation transcriptional programmes. To explore the mechanisms underlying TWIST1-driven migration and proliferation, we integrated bulk RNA-seq data from HAECs and our scRNAseq data from plaque endothelium. This integrative approach identified 45 genes positively-regulated by TWIST1 across both systems (Fig. 3i). From this set, we prioritized 24 genes with known or putative roles in EMT or EndMT and qRT-PCR confirmed TWIST1-dependent regulation for 10 of them (Fig. S8a). These 10 genes were then subjected to siRNA silencing, which was validated by qRT-PCR (Fig. S8b), to assess their potential effects on migration and proliferation. This screening identified a single gene, PELP1, that regulated both processes, and eight genes that specifically regulated proliferation (Fig. S8c-e, Fig. 4, Fig.5).

**Figure 3.**
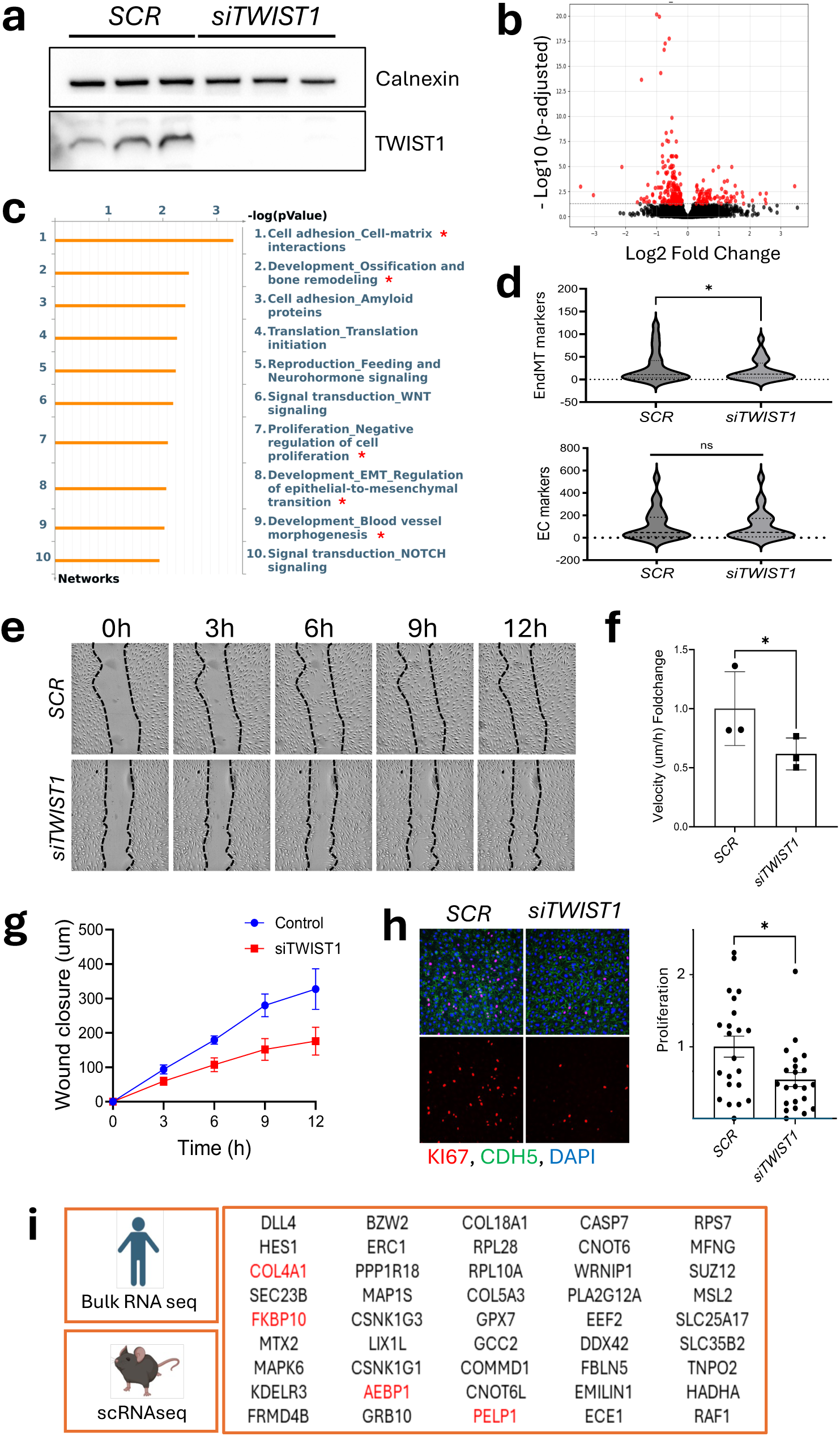
TWIST1 promotes EndMT, proliferation and migration of arterial EC. (A) HAECs were transfected with *TWIST1* siRNA (*siTWIST1*) or scrambled (SCR) siRNA before exposure to DF for 72h (Ibidi system). *TWIST1* knockdown was confirmed by Western blotting (n=3). (B) Volcano plot of RNA-seq data (n= 4) using *SCR* vs *TWIST1*-silenced HAEC under DF using the Ibidi system. Differentially expressed genes (Padj<0.05) are highlighted in red. (C) Process Network Analysis (MetaCore) identified biological processes enriched by *TWIST1 siRNA* in RNA-seq. Pathways related to EndMT or cell proliferation are indicated by red asterisks. (D) Violin plot of EMT-related gene expression (top) and EC-specific markers (bottom) from RNA-seq data in *SCR* vs *siTWIST1* treated HAECs (n=4). (E) Cell migration was assessed using a scratch wound assay in *SCR* vs *siTWIST1* HAEC monolayers after exposure to DF (orbital shaker). Brightfield images show wound closure at different time points (scale bar= 200 μm). (F) Average migration velocity over 12 hours was quantified (distance migrated/12h) (n=3). (G) The distance migrated from the initial wound (T0) was measured at multiple time points. (H) Ki67 immunofluorescence staining (red) was performed to quantify proliferation in *SCR* vs *siTWIST1*-treated HAEC under DF (Ibidi system). Merged images show DAPI (nuclear stain, blue) and VE-cadherin (green) (scale bar= 100 μm). Ki67-positive cells were quantified as a proportion of total nuclei and values are shown as relative fold change compared to *SCR* (n=3). (G) RNA-seq analysis of HAECs was compared with scRNAseq from murine plaque endothelium, identifying 45 common *Twist1*-regulated genes involved in EMT, EndMT, proliferation, and migration. Mean values are shown +/-standard errors. Differences between means were analysed using a ratio paired t-test (F, H) or a Wilcoxon test (D).

**Figure 4.**
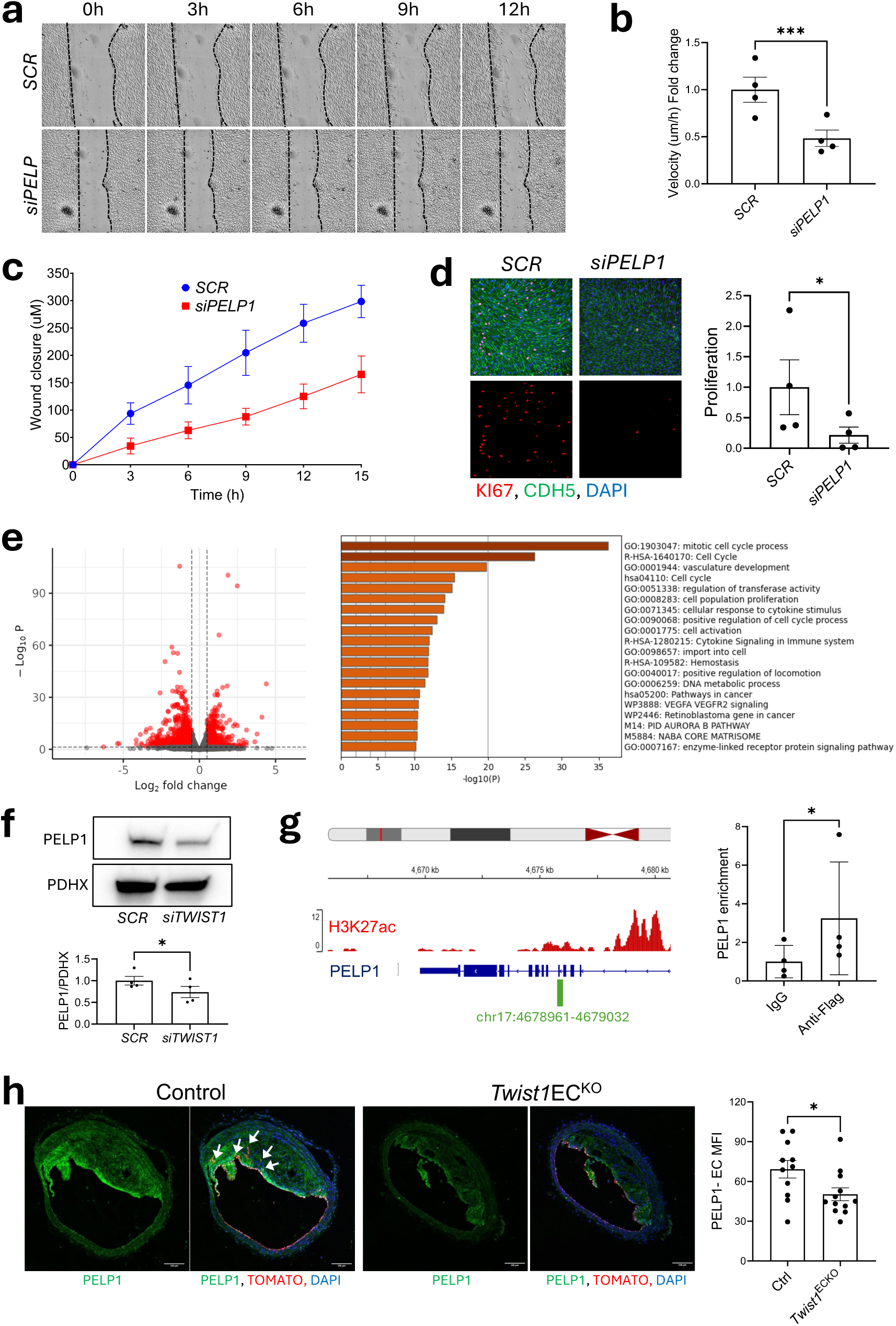
TWIST1 promotes migration and proliferation via PELP1. (A) Cell migration was assessed using a scratch wound assay in *SCR* vs *PELP1*-silenced (*siPELP1*) HAEC monolayers exposed to DF (orbital shaker). Brightfield images show wound closure at different time points (scale bar= 200 μm). (B) Average migration velocity over 12 hours was quantified (distance migrated/12h) (n=4). (C) The distance migrated from the initial wound (T0) was measured at multiple time points. (D) *SCR*-and *siPELP1*-treated HAECs were exposed to DF for 72h (orbital shaker) and stained using antibodies against Ki67 (red) to assess proliferation. Merged images show DAPI (nuclear stain, blue) and VE-cadherin (green) (scale bar= 100 μm). Ki67-positive cells were quantified as a proportion of total nuclei and values are shown as relative fold change compared to *SCR* (n=4). (E) RNA-seq analysis comparing control (*SCR*) and *PELP1*-silenced HAECs exposed to DF using the orbital shaker (n=4). On the left, volcano plot showing differentially expressed genes (Padj<0.05, fold change >1.5, highlighted in red). On the right, pathway enrichment analysis (Metascape) identifying key pathways enriched by *PELP1* siRNA. (F) Western blot analysis of PELP1 expression in *SCR* vs *TWIST1*-silenced HAECs exposed to DF (Ibidi system), normalized to PDHX (n=4). (G) HAECs were infected with lentivirus expressing TWIST1-FLAG and exposed to DF using an orbital shaker for 72h. TWIST1 binding at the *PELP1* loci was assessed using an anti-FLAG antibody, with IgG as a control. On the left, schematic representation of the *PELP1* gene locus, with TWIST1 binding sites highlighted in green. On the right, ChIP-qPCR fold enrichment relative to the IgG control (n=4). (H) Brachiocephalic artery atherosclerotic plaques after 14 weeks of Western diet. Frozen sections of brachiocephalic arteries were stained using antibodies against PELP1 (green) in *Twist1*^ECKO^ (n=12) and control (n=11) mice. Rosa26TdTomato^+^ cells are shown in red and nuclei are counterstained with DAPI (blue). Arrows mark Rosa26TdTomato-PELP1 double-positive cells in the plaque. Representative images are shown (Scale bar=100 μm). Mean values are shown +/-standard errors. Differences between means were analysed using (B, D, F, G) a ratio paired t-test or (H) an unpaired t-test.

**Figure 5.**
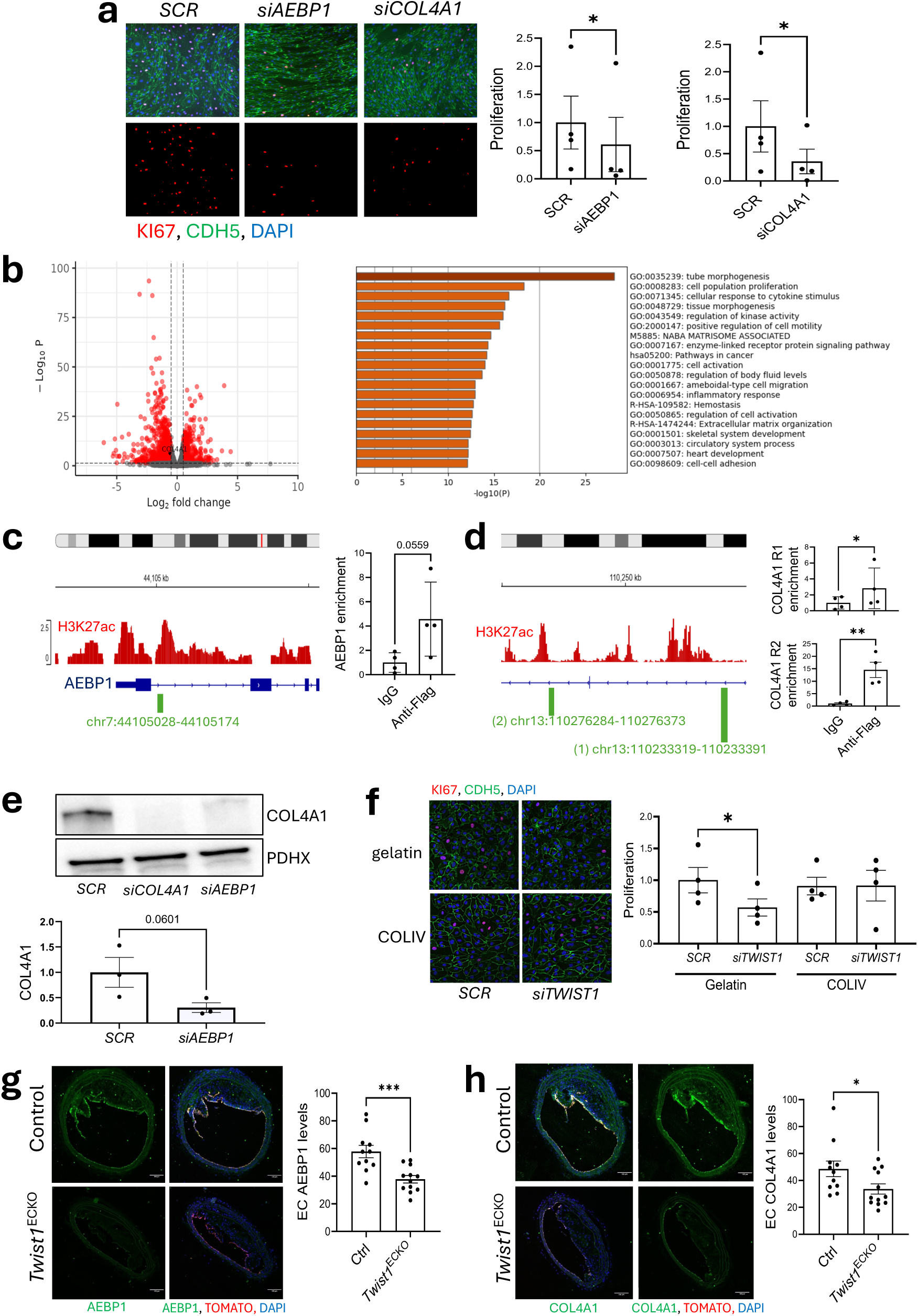
TWIST1 promotes proliferation of arterial EC via AEBP1-COL4A1 signalling. (A) Control (*SCR)*, *AEBP1-*silenced (*siAEBP1),* and *COL4A1*-silenced *(siCOL4A1)* HAECs were exposed to DF (orbital shaker) for 72h, and Ki67 immunofluorescence staining (red) was performed to assess proliferation. Merged images show DAPI (blue) and CDH5 (green) (scale bar= 100 μm). Ki67-positive cells were quantified as a proportion of total nuclei and values are shown as relative fold change compared to *SCR* (n=4). (B) RNA-seq analysis comparing *SCR* and *siAEBP1* HAECs after 72h under DF (orbital shaker). On the left, volcano plot of differentially expressed genes (Padj<0.05, fold change>1.5, highlighted in red). Among these, *COL4A1* is significantly regulated by *AEBP1* and is indicated. On the right, pathway enrichment analysis (Metascape) identifying key pathways enriched by *AEBP1* siRNA. (C, D) HAECs were infected with lentivirus expressing TWIST1-FLAG and exposed to DF using an orbital shaker for 72h. TWIST1 binding at the *AEBP1* and *COL4A1* loci was assessed using an anti-FLAG antibody, with IgG as a control. (C) On the left, schematic representation of the *AEBP1* gene locus, with TWIST1 binding sites highlighted in green. On the right, ChIP-qPCR fold enrichment relative to the IgG control (n=4). (D) On the left, schematic representation of the *COL4A1* gene locus, with TWIST1 binding sites highlighted in green. On the right, ChIP-qPCR fold enrichment relative to the IgG control (n=4). (E) Western blot analysis of COL4A1 levels in control (*SCR)*, *AEBP1-*silenced (*siAEBP1*) and *COL4A1*-silenced (*siCOL4A1*) HAECs (n=3). (F) Proliferation was quantified in *TWIST1*-silenced cells grown on gelatin or a collagen IV matrix by Ki67 immunofluorescence staining (red). Merged images show DAPI (blue) and CDH5 (green) (n=4). Scale bar= 50 μm. (G, H) Brachiocephalic artery atherosclerotic plaques after 14 weeks of Western diet. Frozen sections of brachiocephalic arteries were stained using antibodies against (G) AEBP1 (green) and (H) COL4A1 (green) in *Twist1*^ECKO^ (n=12) and control (n=11) mice. Rosa26TdTomato^+^ cells are shown in red and nuclei are counterstained with DAPI (blue). Representative images are shown (Scale bar=100 μm). Mean values are shown +/-standard errors. Differences between means were analysed using (A, C, D, E) a ratio paired t-test, (F) a one-way anova or (G, H) an unpaired t-test.

Focussing on the transcriptional co-activator PELP1, we observed that it positively regulated both migration (Fig. 4a-4c) and proliferation (Fig. 4d). Consistently, bulk RNA-seq revealed that *PELP1* silencing altered the expression of pathways involved in both of these processes (Fig. 4e). Notably, 34% (99/288) of TWIST1-regulated genes were also regulated by PELP1 (Fig. S9), consistent with a TWIST1-PELP1 pathway. TWIST1 regulation of PELP1 was further validated at the protein level via immunoblotting (Fig. 4f). To determine whether TWIST1 functions as a potential transcriptional activator of *PELP1*, we analyzed the *PELP1* gene. A putative TWIST1 binding sites was identified within a regulatory region marked by H3K27 acetylation (H3K27Ac) (Fig. 4g). ChIP-qPCR on anti-FLAG immunoprecipitates from TWIST1-FLAG-expressing ECs confirmed TWIST1 binding to the PELP1 regulatory region (Fig. 4g), with the SNAI2 promoter as a positive control (Fig. S10), supporting its role in transcriptional regulation. We validated the TWIST1-PELP1 axis *in vivo* by demonstrating that EC-deletion of *Twist1* reduced PELP1 expression in tracked EC within murine brachiocephalic plaques (Fig. 4h). Collectively, these findings suggest that TWIST1 promotes migration and proliferation by upregulating PELP1.

Among the genes regulating proliferation, we focused on the transcription factor AEBP1 and COL4A1, hypothesizing an interaction given the role of AEBP1 as a collagen regulator^24^. Both AEBP1 and COL4A1 positively influenced proliferation (Fig. 5a). Consistently, bulk RNA-seq analysis revealed that AEBP1 silencing affected pathways related to proliferation and matrix remodeling (Fig. 5b). Moreover, comparative RNA-seq analysis identified 23 proliferation modulators co-regulated by AEBP1 and TWIST1 (Fig. S11), supporting a TWIST1-AEBP1 pathway. TWIST1 regulation of AEBP1 and COL4A1 was validated at the protein level (Fig. S12). Consistent with a transcriptional mechanism, ChIP confirmed TWIST1 binding at potential regulatory sites within the AEBP1 and COL4A1 genes (Fig. 5c, 5d). Additionally, siRNA silencing demonstrated that AEBP1 regulates COL4A1 expression (Fig. 5e). At a functional level, replenishing collagen type IV in the extracellular matrix rescued the proliferative capacity of *TWIST1*-silenced cells (Fig. 5f), indicating a critical role for collagen type IV in TWIST1-driven proliferation. We validated our findings *in vivo* by showing that EC-specific *Twist1* deletion reduced AEBP1 (Fig. 5g) and COL4A1 (Fig. 5h) expression in tracked ECs within murine brachiocephalic plaques. Together, these results support a TWIST1-AEBP1-COL4A1 axis that drives EC proliferation.

By integrating data from murine scRNA-seq, cell-based RNA-seq, and gene silencing, we conclude that TWIST1 regulates EndMT in atherosclerosis through a PELP1-mediated migration pathway and an AEBP1-COL4A1-driven proliferation.

### *Twist1* promotes features of plaque stability

To investigate the role of *Twist1* in plaque progression, we analysed disease severity after endothelial cell-specific genetic deletion of *Twist1* in male mice with pre-existing plaques. *Twist1^ECKO^ ApoE^-/-^* mice exhibited a significant reduction in plaque burden compared to controls, suggesting that *Twist1* promotes plaque growth (Fig. 6a). This reduction was not associated with changes in plasma cholesterol or triglyceride levels, which were similar between experimental and control groups (Fig. S13). Notably, plaques in *Twist1^ECKO^ ApoE^-/-^* mice showed reduced fibrillar collagen content (Fig. 6b) and a lower proportion of ACTA2-positive cells (Fig. 6c), both indicators of plaque stability. In contrast, these plaques exhibited increased macrophage content (Fig. 6d), enhanced necrosis (Fig. 6e) and more elastin breaks (Fig. 6f), all features of plaque instability. These findings support the conclusion that *Twist1* plays a critical role in promoting plaque growth while also contributing to features of plaque stability.

**Figure 6.**
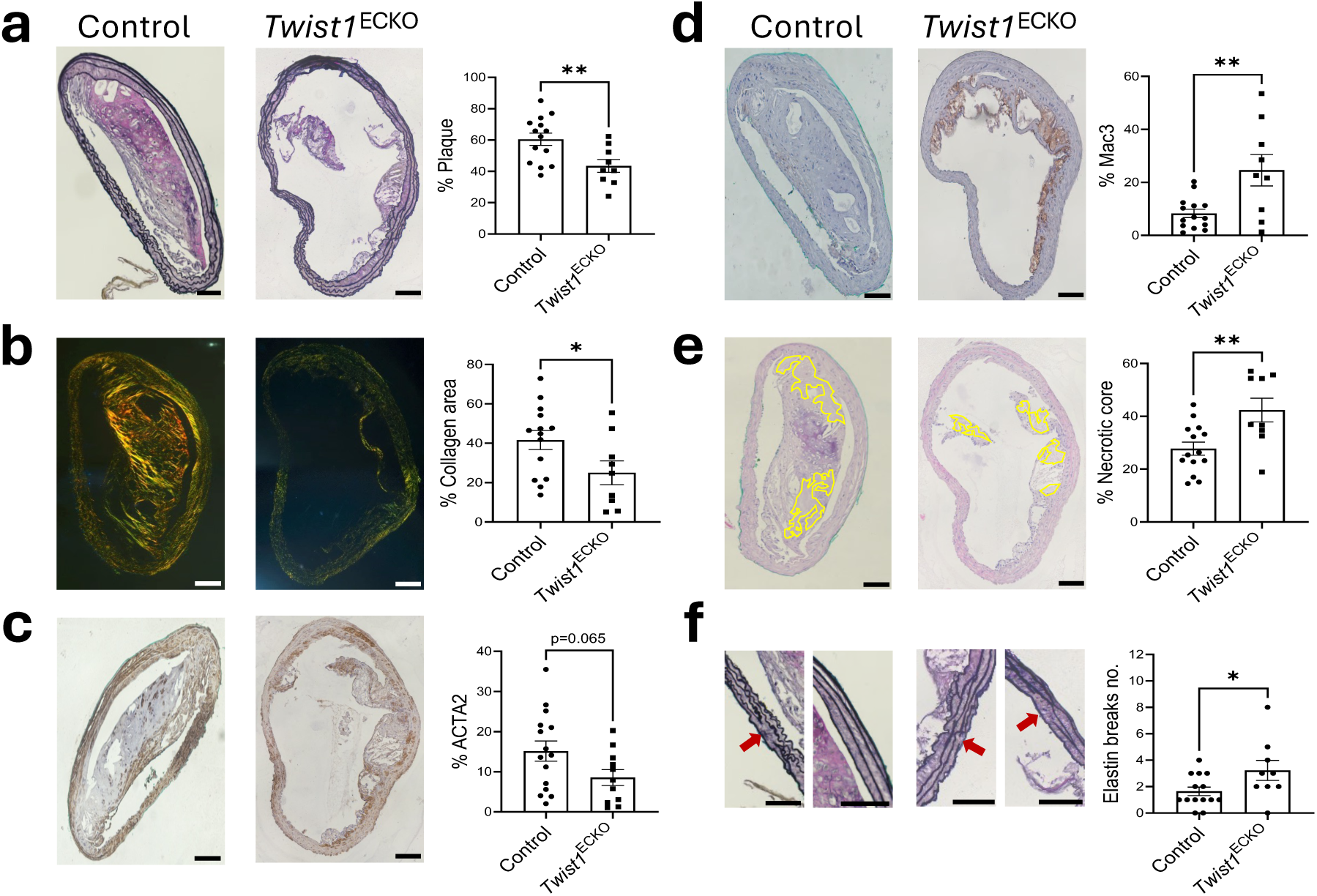
***Twist1* promotes plaque growth and features of stability *in vivo***. Male *Twist1*^ECKO^ (*Twist1^fl/fl^ Cdh5^CreERT^*^2^*^/+^ ApoE^-/-^*) and control mice (*Twist1^fl/fl^ Cdh5^+/+^ ApoE^-/-^*) aged 8 weeks were fed a Western diet for 8 weeks to induce atherosclerotic lesions. Tamoxifen was then administered for 5 consecutive days to induce *Twist1* deletion in ECs and a Western diet was provided for an additional 6 weeks (totalling 14 weeks of Western diet). Paraffin-embedded sections of brachiocephalic arteries were stained with (A) Miller’s elastin stain to quantify plaque burden, (B) Picrosirius Red (visualised under polarised light) to quantify collagen content, (C) antibodies against ACTA2 to quantify vSMCs content, (D) antibodies against MAC3 to quantify macrophage content, and (E) Hematoxylin and Eosin (H&E) to quantify necrotic core (highlighted in yellow) content in *Twist1*^ECKO^ (n=9) and control (n=14) mice. (F) Magnified view of specific regions shown in (A) and quantification of elastin breaks number in brachiocephalic arteries from *Twist1*^ECKO^ (n=9-11) and control (n=14-15) mice. The red arrows indicate elastin breaks. Representative images are shown (Scale bar=100 μm). Mean values are shown +/-standard errors. Differences between means were analysed using an unpaired t-test.

### Sex differences in *Twist1* control of EndMT and features of stability

We conducted parallel experiments to analyse the influence of *Twist1* on EndMT and plaque stability in female mice. EC tracking demonstrated EndMT in brachiocephalic plaques of female mice (Fig. S14a). However, endothelial deletion of *Twist1* did not influence EndMT (Fig. S14a), plaque burden (Fig. S14b), or features of plaque stability (Fig. S14c-g) in female mice. These findings indicate that *Twist1* regulation of EndMT and its impact on features of plaque stability are observed in male mice but not in females, highlighting the sex-dependent mechanisms of atherosclerosis.

### TWIST1 predicts stabilization of human carotid artery plaques

To translate these findings to human disease, we quantified the expression of TWIST1 in human carotid plaques from the Carotid Plaque Imaging Project (CPIP, Malmö, Sweden) biobank. Patients were classified as symptomatic if they had experienced amaurosis fugax, transient ischemic attack, or ischemic stroke within one month prior to surgery. Bulk RNAseq from the most stenotic region of the plaque showed significantly higher TWIST1 mRNA levels in asymptomatic plaques compared to symptomatic plaques (Fig. 7a). A key strength of the Malmö cohort is its longitudinal follow-up, allowing the analysis of markers that may predict future cardiovascular events. Notably, higher TWIST1 expression (2^nd^ and 3^rd^ tertile) was associated with a reduced risk of future cardiovascular events during post-operative follow-up period of up to five years compared to low levels (1^st^ tertile; Fig. 7b). Finally, we identified TWIST1 protein expression in human carotid plaque endothelial and vascular smooth muscle cells using immunofluorescence (Fig. 7c). In conclusion, TWIST1 is associated with plaque stability and serves as a predictive marker of favourable cardiovascular outcomes in patients after carotid endarterectomy.

**Figure 7.**
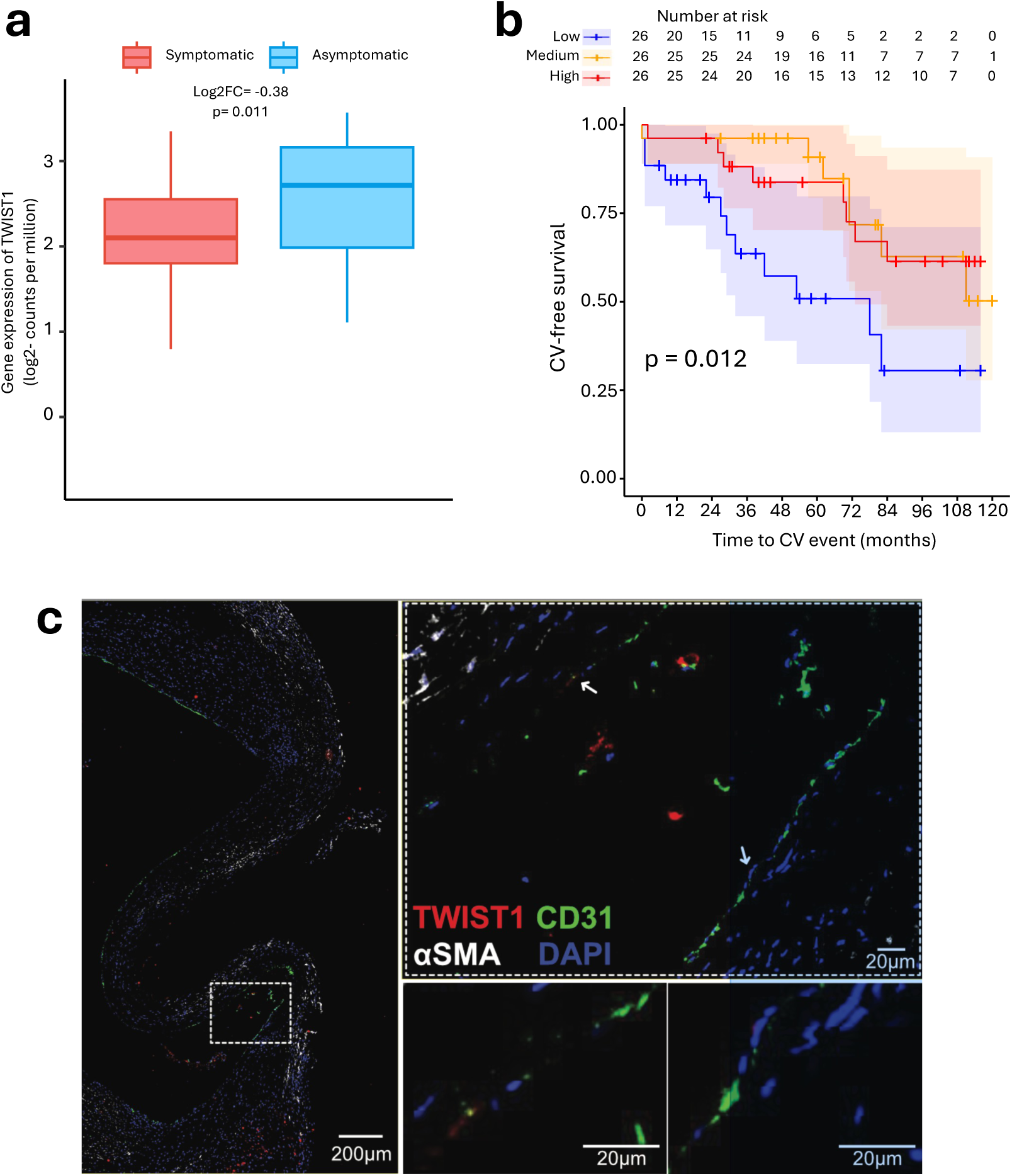
*TWIST1* expression in human carotid plaques is associated with asymptomatic disease and favourable outcome. (A, B) *TWIST1* expression levels in human carotid artery endarterectomies were analyzed using bulk RNA sequencing. (A) *TWIST1* levels were significantly lower in plaques from symptomatic patients (n=51) compared to asymptomatic ones (n=27). (B) Kaplan–Meier survival curves showing that lower plaque *TWIST1* mRNA levels (1^st^ tertile compared to 2^nd^-3^rd^ tertile of *TWIST1* mRNA levels) predict post-operative cardiovascular events. (C) Representative images of fluorescent staining for TWIST1 (red), the EC marker CD31 (green), and the vSMC markers αSMA (white) with nuclei counterstained using DAPI (blue) (n = 3). The right panel shows a magnified view of the region highlighted by the white box. Arrows indicate TWIST1 expression in ECs.

## DISCUSSION

EndMT contributes to the cellular composition of advanced atherosclerotic plaques ^10,11,13–17^. Studies in hypercholesterolemic mice have shown that modulating EndMT-related pathways influences disease progression^10,11,15–17^, further supporting the notion that EndMT is commensurate with disease progression. Similarly, analyses of atherosclerotic plaques in human coronary arteries^10^ and the abdominal aorta^13^ have revealed increased EndMT marker expression in advanced disease compared to earlier stages. These observations have led to the suggestion that EndMT plays a pathogenic role in the progression of atherosclerosis towards unstable rupture-prone lesions. Our findings challenge this perspective, demonstrating that *Twist1* promotes EndMT while increasing plaque stability. Although this conclusion goes against the current paradigm, it is nevertheless intuitive because several canonical features of EndMT, such as collagen deposition and the accumulation of ACTA2-positive cells, are also hallmarks of plaque stability.

To integrate our findings with existing literature, we propose that EndMT in atherosclerotic plaques represents a phenotypic continuum with diverse functional outcomes. Our scRNA-seq analysis supports this interpretation, identifying four clusters enriched in EndMT markers. Among them, Cluster 8 exhibited a more advanced EndMT phenotype, characterized by the downregulation of endothelial markers alongside the upregulation of EndMT markers. In contrast, Clusters 2, 3, and 7 retained endothelial markers alongside EndMT markers, indicative of partial EndMT—a phenomenon previously described in angiogenesis ^25,26^, pulmonary hypertension^20,27^, and post-myocardial infarction remodeling^28^. TWIST1 promoted Clusters 2, 7, and 8 but had no effect on Cluster 3, suggesting that TWIST1 regulates specific subsets of EndMT cells in atherosclerosis. This idea of multiple EndMT subsets aligns with several scRNA-seq analyses that have underscored the phenotypic heterogeneity of ECs across organs and disease states, revealing that EndMT represents a spectrum of phenotypes rather than a singular state ^29^ ^12,30^. It also resonates with observations of EMT, where epithelial cells transition into various mesenchymal states, leading to significant heterogeneity in cell fates and functions^31–33^. It is likely that different forms of EndMT and EMT are controlled by transcription factors that exert distinct roles^32,33^. For example, Snai1 and Prrx1 drive different modes of EMT during vertebrate development^32^. By analogy, our data suggest that TWIST1 regulates protective forms of EndMT in advanced atherosclerosis, and we propose that other unidentified factors contribute to the pathogenic forms of EndMT. Further research is needed to test this hypothesis and better understand the functional diversity of EndMT in atherosclerosis.

In this study, we delineated the role of TWIST1 in driving EndMT within atherosclerotic plaques, identifying its critical involvement in migration and proliferation pathways. TWIST1 achieves this by inducing the expression of multiple genes, with our analysis focusing on PELP1 and AEBP1. We found that PELP1 is required for both migration and proliferation, while AEBP1 is specifically required for proliferation. The involvement of PELP1 in EndMT is consistent with its known role in EMT and subsequent cell migration in breast cancer^34,35^. Our findings reveal a novel role for AEBP1 in EndMT, aligning with its established involvement in EMT^31^ and epithelial cell proliferation ^36^. AEBP1 acts as a central regulator of collagen, with mutations in the AEBP1 gene linked to defective collagen assembly in Ehlers-Danlos syndrome^24^. Additionally, AEBP1 has been implicated in regulating cell plasticity and activation in fibrosis^37,38^, as well as upregulating collagen production^39,40^. We demonstrated that AEBP1 induces endothelial expression of COL4A1, a basement membrane component critical for vascular homeostasis^41–43^ and cell proliferation^44,45^. Indeed, COL4A1 mutations are associated with coronary artery disease^46^ and stroke^47^. Our findings suggest that COL4A1 promotes the proliferation of EndMT cells downstream of the TWIST1-AEBP1 pathway. Further research is needed to determine whether COL4A1 contributes to plaque stability through this mechanism. Overall, TWIST1 regulates EndMT via PELP1 and AEBP1-COL4A1 signaling, promoting enhanced migration and proliferation.

*Twist1*-driven EndMT and plaque stability were observed in male but not female mice, highlighting sex-specific differences in atherosclerosis mechanisms. In women, atherosclerosis often presents as diffuse involvement of the cardiovascular system, including microvascular disease, and is associated with a more stable plaque phenotype. In contrast, men typically develop focal disease with larger, more vulnerable plaques^48,49^. Notably, plaque rupture and erosion, key triggers of stroke, unstable angina, and myocardial infarction, exhibit sex-specific patterns, with plaque erosion most common in pre-menopausal women^48,49^. Our observation that atherosclerosis is unaltered in female *Twist1^ECKO^* mice is consistent with Shin *et al* who demonstrated that EndMT is more prominent in atherosclerotic lesions of male *ApoE^-/-^* mice compared to females^50^. These findings highlight the need for further research into the sex-dependent role of EndMT in atherosclerosis.

In summary, we identify TWIST1 as a critical regulator of specific clusters of EndMT cells in advanced atherosclerosis. TWIST1-dependent EndMT enhances endothelial migration via PELP1 and activates an AEBP1-COL4A1 pathway to drive endothelial proliferation. Our findings challenge the prevailing view that EndMT solely destabilizes plaques, instead highlighting a TWIST1-driven program that promotes plaque stability to reduce rupture risk. Further research is necessary to unravel the molecular regulation of EndMT subtypes in atherosclerosis and assess the potential of TWIST1 as a therapeutic target for stabilizing vulnerable plaques.

## METHODS

### Mice

To generate *Twist1^fl/fl^ Cdh5^CreERT^*^2^*^/+^ ApoE^-/-^* mice (*Twist1*^ECKO^ ApoE^-/-^ and control mice), *Twist1^fl/fl^ CDH5^Cre-ERT^*^2^ mice were crossed with *ApoE ^-/-^*mice. To generate *Twist1^fl/fl^ Cdh5^CreERT^*^2^*^/+^ ApoE^-/-^ Rosa26^TdTomato/TdTomato^* mice (*Twist1*^ECKO^ ApoE^-/-^ and control mice on a EC-fate mapped background), *Twist1^fl/fl^ CDH5^Cre-ERT^*^2^ *Rosa26^tdTomato/tdTomato^* mice were crossed with *ApoE ^-/-^*mice. All mice were on a C57BL/6J background. Genotyping was performed using PCR primers described in Table S1. Mice aged 8 weeks old were given a Western diet (TD.88137, Envigo) for 8 weeks. Cre was then activated by intraperitoneal administration of tamoxifen (Sigma-Aldrich) in corn oil for five consecutive days (2 mg per mouse per day), followed by a high-fat diet for 6 weeks. C57BL/6 and transgenic mice were housed in ventilated cages. Animal care and experimental procedures were carried out under licenses issued by the UK Home Office, and local ethical committee approval was obtained.

### Immunohistochemical staining of murine tissues

Mice were euthanized and perfusion-fixed with phosphate-buffered saline (PBS), followed by 4% paraformaldehyde (PFA) via the left ventricle of the heart. The brachiocephalic artery was dissected, gently cleaned of adventitial tissue, and processed for paraffin or frozen immunohistochemical staining. For paraffin sections, blocks were sectioned at 5 μm intervals. Slides were dewaxed in xylene and processed for staining with antibodies (Anti-ACTA2, anti-Mac3, (see Table S2), hematoxylin-eosin staining, Miller’s elastin staining and Picrosirius staining. Samples were boiled for 10 mins in citrate buffer for antigen retrieval. Sections were imaged using a Leica Thunder microscope and analysed using ImageJ. For frozen sections, brachiocephalic arteries were placed in 30% sucrose for 72 hours before embedding in optical cutting temperature compound and snap-freezing. Blocks were sectioned at 10 μm intervals. After washing with PBS and permeabilising with 0.5% Triton X-100 for 20 mins, slides were blocked with 1% bovine serum albumin (BSA)s, 10% horse serum and stained with primary antibodies (anti-VWF, anti-ACTA2, anti-COL4A1, anti-AEBP1 or anti-PELP1, (see Table S2) overnight at 4°C. Sections were then washed with PBS, incubated with Alexa Fluor 488-conjugated secondary antibodies for 1hr at room temperature, washed again with PBS, incubated with DAPI and mounted with ProLong Gold mounting reagent. Frozen sections were imaged using a Zeiss LSM 980 Airyscan confocal microscope and analysed using ImageJ.

### Plasma Lipid Measurements

Blood samples collected by terminal cardiac puncture were subjected to centrifugation to obtain plasma. Plasma cholesterol and triglyceride levels were measured using a COBAS analyser (total plasma cholesterol, non–high-density lipoprotein cholesterol, high-density lipoprotein cholesterol, low-density lipoprotein cholesterol and triglycerides).

### Mouse aorta digestion and FACS for scRNA-seq

The left ventricle of the murine heart was perfused with heparin sodium (20 U/ml) in PBS. Aortas from male *Twist1*^ECKO^ (*Twist1^fl/fl^ Cdh5^CreERT^*^2^*^/+^ ApoE^-/-^*) (n=5) and control (*Twist1^fl/fl^ Cdh5^+/+^ ApoE^-/-^*) (n=4) mice after 14 weeks of Western diet were dissected from the aortic root to the iliac artery, removing fat and connective tissue. Aortas were then transferred into PBS and incubated in collagenase type I (450 U/ml), collagenase type XI (125 U/ml), hyaluronidase type 1-s (60 U/ml), deoxyribonuclease I (60 U/ml), and elastase (0.5 mg/ml) in PBS for 10 min at 37°C, to enable the removal of the adventitial layer. Aortas were then cut into 2 mm pieces and further incubated in the enzymatic solution for 1 hour and 15 mins at 37°C to generate a single-cell suspension. When the aortas were digested, the single-cell suspensions were filtered through a 40-μm cell strainer, washed, and resuspended in 1% BSA solution in PBS. Single-cell suspensions were incubated for 5 to 10 min with TruStain FcX (anti-mouse CD16/32) antibody to block for unspecific binding of immunoglobulin to Fc receptors, and they were subsequently stained with allophycocyanin (APC)-conjugated anti-CD45 and Alexa Fluor (AF) 488–conjugated anti-CD31, washed and diluted in 1% BSA in PBS. To exclude dead cells, samples were stained with TOPRO-3. CD31^+^, CD45^−^, and TOPRO-3^−^ cells were sorted into 384-well plates containing reverse transcription primers, deoxyribonucleotide triphosphates, External RNA Controls Consortium (ERCC) RNA Spike-Ins, and mineral oil using a BDFACSMelody cell sorter (BD Biosciences). Plates containing capture cells were snap-frozen and stored at −80°C until sequencing was peformed.

### Single-cell RNA sequencing

scRNA-seq libraries were generated using the SORT-seq approach as described previously^51^. After lysing single cells at 65°C for 5 min, reverse transcriptase and second-strand mixers were added to the wells using the Nanodrop II liquid handling platform (GC Biotech). The mRNA of each individual cell was reverse-transcribed, double-stranded complementary DNAs were pooled, and *in vitro* transcription was carried out for linear amplification of RNA. The Illumina sequencing libraries were generated using TruSeq small RNA primers (Illumina), and the Illumina NextSeq (carried out commercially by Single Cell Discoveries, Utrecht, The Netherlands) was used to sequence paired-end at 75–base pair read length the DNA libraries. The RNA yield of the amplified RNA and the quality and concentration of the final cDNA libraries were measured using Bioanalyzer (Agilent).

### Bioinformatics

Reads from Illumina sequencing were aligned to the GRCm38 mouse genome using STAR. Single-cell transcriptomes generated using SORT-seq were quality-filtered, excluding cells with less than 1800 transcripts per cell and more than 10% mitochondrial content from further analysis (2139 cells out of 3043 total sorted cells were analysed). After filtering count data and selecting high-quality single cells, the data were normalized, and the top 2000 highly variable genes in our dataset were selected and used for principal component analysis. The first 20 principal components were retained based on their contribution to overall variability and used for downstream analysis, to perform dimensional reduction and clustering. Dimensional reduction and clustering of our scRNA-seq dataset were performed using the R package Seurat (v3.1) and a clustering resolution of 0.8 was used. Cell cycle regression was performed by calculating an aggregated score for S and G2M genes per cell to mitigate the effects of cell cycle heterogeneity. Differential expression analysis was performed between clusters, and marker genes for each cluster were determined with the Wilcoxon rank sum test with *P* < 0.001 using Seurat and the single-cell analysis platform BBrowserX. Differential expression analysis, heatmaps of gene expression embedded on hierarchical clustering, t-distributed stochastic neighbour embedding (t-SNE) representations showing the expression of defined genesets/GO pathways, and expression of single transcripts on the t-SNE embedding were performed using the software BBrowserX. Violin plot representations of the expression of defined GO pathways shows the AUCellScore, which uses the “Area Under the Curve” to calculate whether a critical subset of selected GO pathways is enriched within the expressed genes of each cell.

### Cell culture and flow experiments

HAECs were obtained from PromoCell and cultured in EGM2 growth medium (#C-22011) according to the manufacturer’s guidelines. HAECs from four different donors (three male, one female), non-pooled, were used for all experiments between passages 3 and 7. Cells were exposed to flow conditions using either the Ibidi flow apparatus or an orbital shaker. For Ibidi flow, cells were seeded on gelatin or collagen IV-coated Ibidi μ-Slides I^0.4^ (Luer ibiTreat, Ibidi) and used when cells reached full confluence. UF consisted of 13 dynes/cm² unidirectional shear stress. To generate DF, HAECs were subjected to a cyclic regimen of 2 hours of oscillatory flow (±4 dynes/cm², 0.5 Hz), followed by 10 minutes of unidirectional flow (+4 dynes/cm²) to promote nutrient redistribution. The slides and pump system were maintained in a cell culture incubator at 37°C during the experiment. For orbital flow, HAECs were grown to confluence in gelatin-coated 6-well plates and exposed to flow for 72 hours on an orbital shaking platform (PSU-10i; Grant Instruments) placed inside a cell culture incubator. The shaker had a 10 mm orbit radius and was set at 210 revolutions per minute (RPM), generating DF (approximately 5 dynes/cm² multidirectional shear stress) at the well center and uniform flow with higher shear stress (approximately 10 dynes/cm²) and stable directionality at the periphery.

### Gene silencing

HAECs were transfected with siRNA sequences targeting TWIST1 (L-006434-00, Dharmacon), AEBP1 (s1145, Ambion), COL4A1 (s3289, Ambion), DLL4 (s29215, Ambion) FKBP10 (s34173, Ambion), KDELR3 (s21690, Ambion), MTX2 (s20935, Ambion), RPS7 (s12290, Ambion), (s34173, Ambion), PELP1 (s25746, Ambion), SEC23B (s20536, Ambion), USP14 (s17358, Ambion) using the Lipofectamine RNAiMAX transfection system (13778-150, Invitrogen), as per the manufacturer’s protocol. Scrambled siRNA (D-001810-01-50, Dharmacon) was used as a control. Experiments were conducted with 25 nM siRNA, and cells were incubated in transfection media for 6 hours before media replacement and exposure to flow.

### Quantitative Real-Time PCR (qRT-PCR)

Total RNA was extracted from HAECs using the RNeasy Mini Kit (Qiagen, 74104) and reverse transcribed into cDNA using the iScript cDNA Synthesis Kit (Bio-Rad, 1708891). Gene expression was measured by qRT-PCR with specific primers (Table S2) and SsoAdvanced Universal SYBR Green Supermix (Bio-Rad, 172-5271). Expression levels were normalized to the housekeeping gene HPRT, and fold changes were calculated using the ΔΔCt method.

### Bulk RNA sequencing of cultured EC

Total RNA from HAECs was extracted and quality-checked using a Bioanalyzer (Agilent). High-quality RNA was used for RNA-seq library preparation and sequenced on an Illumina platform, generating 40 million reads per sample. Library preparation and sequencing were performed by Novogene. Fastq files were processed using the nf.core rna-seq pipeline (version 3.6.0) (https://nf-co.re/rnaseq/3.6) incorporating salmon pseudo-alignment and quantification. The DESeq2 Bioconductor package was used for differential expression, with a donor effect added to the model to compensate for donor-specific effects in expression observed with a PCA plot. Functional enrichment analysis was conducted using Metacore/Metascape on protein-coding genes with P < 0.05.

### Western Blot

Flow-exposed cells were washed twice with PBS and lysed in Laemli/RIPA buffer buffer containing β-mercaptoethanol. Lysates were separated on 4–15% Mini-PROTEAN® TGX™ Precast Protein Gels, (Biorad #4561083) for 90 minutes at 100V, followed by transfer to PVDF membranes (Bio-Rad, #1660827 EDU) at 230 mA for 60 minutes. Membranes were blocked for 1 hour with 10% milk in TBS-T, then stained overnight with primary antibodies (Table S3) in 5% milk. After washing, secondary antibodies (Goat Anti-Mouse or Goat Anti-Rabbit, Dako 0447 and P0448) were applied at 1:3000 in 5% milk for 1 hour. Blots were visualized using ECL Select (Amersham, #RPN2235) and the Chemidoc XRS system (Bio-Rad).

### Immunofluorescence of cultured EC

HAECs were washed twice with PBS, fixed for 10 minutes in 4% paraformaldehyde, permeabilized with 0.1% Triton X-100, and blocked with 20% goat serum. Cells were incubated with primary antibodies (Table S3) in 5% serum overnight at 4°C. After washing, secondary antibodies conjugated with Alexa Fluor were applied, and images were captured using a Leica Thunder, LSM980 Confocal, or ZEISS CD7 microscope.

### Migration assay

HAECs grown on gelatin-coated 6-well plates and silenced for TWIST1, PELP1, or scrambled siRNA were exposed to 72 hours of orbital flow-induced DF. Cells were scratched using a pipette tip, washed, and fresh media was added. Wound closure was monitored using live imaging on the ZEISS CD7 microscope at 0, 3, 6, 9 and 12 hours post-incubation. Migration rates were quantified by measuring the distance moved by cells from the scratch edge. The velocity was calculated as the distance/hour after 12hours assay.

### Chromatin Immunoprecipitation (ChIP)

ChIP assays were performed using the Diagenode iDeal ChIP qPCR kit, as per the manufacturer’s instructions. HAECs overexpressing TWIST1-Flag were subjected to 72 hours of orbital flow before cross-linking with 1% formaldehyde. Cells exposed to DF were scratched and lysed to isolate nuclei. chromatin was sheared via sonication 30sec ON/ 30sec OFF on a bioruptor plus sonicator (Diagenode). Immunoprecipitation was performed using protein A magnetic beads and 10 μg of ChIP grade anti-Flag or rabbit IgG antibody for 3 hours, followed by overnight incubation with sheared chromatin. Immunoprecipitated DNA was quantified using qPCR with primers listed in (Table S2). DNA enrichment over input was calculated, and the data were normalized to IgG.

### TWIST1 Overexpression

To overexpress TWIST1, HEK 293T cells were plated on Day 1 to reach 80-90% confluency by the following day. On Day 2, cells were transfected with a plasmid mix containing 4,2pm of Pax2, VSVG, and Twist1-Flag vector using PEI (polyethylenimine). The plasmid-PEI complex was incubated for 20 minutes before being added to the cells. After 6 hours, the media was replaced with MEM (+10% FCS, glutamin, Pen/Strep). On Day 3, media was changed to EGM2. Viral supernatants were collected on Days 4 and 5, centrifuged at 500g, filtered through a 0.45 μm filter, and transferred to HAECs at a 1:1 ratio.

### Human carotid plaque clinical data, bulk RNA-sequencing and survival analysis

Human plaque bulk RNAseq and immunoflouresent staining of TWIST1 and CD31 were performed on carotid plaques obtained from the CPIP biobank (Region Skåne, Malmö, Sweden). All patients have given written informed consent, and the study follows the declaration of Helsinki. The study protocol has been approved by the local ethical committee in Lund and the Swedish ethical committee (472/2005, 2014/904, 60/2008, 2012/209). The indications for surgery were: 1) asymptomatic carotid stenosis with a stenosis degree >80% or 2) cerebrovascular symptoms (<u>ischemic</u> <u>stroke</u>, <u>transient ischemic attack</u>, or <u>amaurosis fugax</u> within one month prior to surgery) and a carotid plaque with stenosis degree >70%, as previously described^52^.

All plaques were immediately snap frozen in liquid nitrogen upon surgical removal. To assess TWIST1 mRNA levels one cross-sectional fragment of 1mm was taken from the most stenotic part of all plaques. RNA was isolated using Trizol and sequenced using both Illumina HiSeq2000 and NextSeq 500 platforms, as described previously^53^. In short, transcript-level quantification was conducted using Salmon based on transcriptome release 27 of GENCODE in mapping-based mode. Gene counts were summarized using tximport and were normalized between samples using a trimmed mean of M-values (TMM) by edgeR^54^, giving gene expressions as log2-transformed counts per million (CPM) after voom transformation. Batch effects of sequencing platforms were adjusted by an empirical Bayes method^55^. For follow-up analysis, information regarding postoperative cardiovascular events (myocardial infarction, unstable angina, stroke (ipsilateral and contralateral events), transient ischemic attack, amaurosis fugax, vascular interventions (including carotid endarterectomy/stenting, coronary artery bypass grafting/percutaneous coronary artery intervention) and CV death was acquired from the Swedish Cause of Death and National inpatient Health Registers. The participants were followed until events or end of follow-up by the 31^st^ December 2015.

### Multiplex immunofluorescence staining of human atherosclerotic plaques

Formalin-fixed, decalcified and paraffin-embedded human atherosclerotic plaque tissue sections (4μm) were stained using antibodies to CD31, TWIST1 and ACTA2, and visualized with the corresponding Opal fluorophores 520, 620 and 780 (1:100 dilution), following Akoya’s Phenoptics workflow. In brief, sections underwent dewaxing, rehydration, and antigen retrieval in citric buffer (pH 6) prior to staining. Each staining cycle included primary antibody incubation, HRP-conjugated secondary application, tyramide signal amplification with Opal fluorophores, and stripping using β-mercaptoethanol-based buffer (50°C for 30 min) to remove antibodies while retaining Opal-conjugated signals. A spectral library for multiplex slide analysis was prepared, including single-stained control slides, a slide with only DAPI, and an autofluorescence control slide. Multispectral imaging was conducted using PhenoImager HT (Akoya Biosciences) and analyzed with inForm software and Qupath.

### Statistical analysis

Data are presented as mean values ± standard error of the mean. GraphPad Prism analysis software was used to carry out statistical analyses. Significance is shown as follows: *p<0.05; **p<0.01; ***p< 0.001, ****p<0.0001. Specific tests used for each experiment are explained in the figure legends.

## DATA AVAILABILITY

The fully annotated scRNA-seq and bulk RNA-seq datasets are publicly accessible at the Gene Expression Omnibus [GSE293412]. The human data is protected due to privacy laws and would be shared in group form upon request from a qualified academic investigator for the sole purpose of replicating the procedures and results presented in the article and providing that the data transfer is in agreement with European Union legislation on the general data protection regulation and decisions by the ethical review board of Sweden, the Region Skåne and the Lund University. Professor Isabel Goncalves may be contacted for data access. Data regarding living subjects cannot be publicly available due to the sensitive nature of the data regulated by GDPR.

## ACKNOWLEDGEMENTS

We acknowledge Lena Sundius and Fiona Wright for their excellent histology support.

## REFERENCES

1. Buono MF, Slenders L, Wesseling M, Hartman RJG, Monaco C, den Ruijter HM, Pasterkamp G, Mokry M. The changing landscape of the vulnerable plaque: a call for fine-tuning of preclinical models. Vascul Pharmacol. 2021;141:106924. doi: 10.1016/j.vph.2021.106924

2. Hoogendoorn A, Kok AM, Hartman EMJ, de Nisco G, Casadonte L, Chiastra C, Coenen A, Korteland SA, Van der Heiden K, Gijsen FJH, et al. Multidirectional wall shear stress promotes advanced coronary plaque development: comparing five shear stress metrics. Cardiovasc Res. 2020;116:1136–1146. doi: 10.1093/cvr/cvz212

3. Frid MG, Kale VA, Stenmark KR. Mature vascular endothelium can give rise to smooth muscle cells via endothelial-mesenchymal transdifferentiation: in vitro analysis. Circ Res. 2002;90:1189–1196. doi: 10.1161/01.res.0000021432.70309.28

4. Souilhol C, Harmsen MC, Evans PC, Krenning G. Endothelial-mesenchymal transition in atherosclerosis. Cardiovascular Research. 2018;114:565–577. doi: 10.1093/cvr/cvx253

5. Hall IF, Kishta F, Xu Y, Baker AH, Kovacic JC. Endothelial to mesenchymal transition: at the axis of cardiovascular health and disease. Cardiovasc Res. 2024;120:223–236. doi: 10.1093/cvr/cvae021

6. Nakajima Y, Yamagishi T, Hokari S, Nakamura H. Mechanisms involved in valvuloseptal endocardial cushion formation in early cardiogenesis: roles of transforming growth factor (TGF)-beta and bone morphogenetic protein (BMP). Anat Rec. 2000;258:119–127. doi: 10.1002/(sici)1097-0185(20000201)258:2<119::Aid-ar1>3.0.Co;2-u

7. DeRuiter MC, Poelmann RE, VanMunsteren JC, Mironov V, Markwald RR, Gittenberger-de Groot AC. Embryonic endothelial cells transdifferentiate into mesenchymal cells expressing smooth muscle actins in vivo and in vitro. Circ Res. 1997;80:444–451. doi: 10.1161/01.res.80.4.444

8. Moonen JA, Lee ES, Schmidt M, Maleszewska M, Koerts JA, Brouwer LA, Van Kooten TG, Van Luyn MJ, Zeebregts CJ, Krenning G, et al. Endothelial-to-mesenchymal transition contributes to fibro-proliferative vascular disease and is modulated by fluid shear stress. Cardiovascular research. 2015. doi: 10.1093/cvr/cvv175

9. Mahmoud MM, Kim HR, Xing RY, Hsiao S, Mammoto A, Chen J, Serbanovic-Canic J, Feng S, Bowden NP, Maguire R, et al. TWIST1 Integrates Endothelial Responses to Flow in Vascular Dysfunction and Atherosclerosis. Circulation Research. 2016;119:450-+. doi: 10.1161/circresaha.116.308870

10. Chen PY, Qin L, Baeyens N, Li G, Afolabi T, Budatha M, Tellides G, Schwartz MA, Simons M. Endothelial-to-mesenchymal transition drives atherosclerosis progression. J Clin Invest. 2015;125:4514–4528. doi: 10.1172/jci82719

11. Kidder E, Pea M, Cheng S, Koppada SP, Visvanathan S, Henderson Q, Thuzar M, Yu X, Alfaidi M. The interleukin-1 receptor type-1 in disturbed flow-induced endothelial mesenchymal activation. Front Cardiovasc Med. 2023;10:1190460. doi: 10.3389/fcvm.2023.1190460

12. Andueza A, Kumar S, Kim J, Kang DW, Mumme HL, Perez JI, Villa-Roel N, Jo H. Endothelial Reprogramming by Disturbed Flow Revealed by Single-Cell RNA and Chromatin Accessibility Study. Cell Rep. 2020;33:108491. doi: 10.1016/j.celrep.2020.108491

13. Evrard SM, Lecce L, Michelis KC, Nomura-Kitabayashi A, Pandey G, Purushothaman KR, d’Escamard V, Li JR, Hadri L, Fujitani K, et al. Endothelial to mesenchymal transition is common in atherosclerotic lesions and is associated with plaque instability. Nat Commun. 2016;7:11853. doi: 10.1038/ncomms11853

14. Newman AAC, Serbulea V, Baylis RA, Shankman LS, Bradley X, Alencar GF, Owsiany K, Deaton RA, Karnewar S, Shamsuzzaman S, et al. Multiple cell types contribute to the atherosclerotic lesion fibrous cap by PDGFRβ and bioenergetic mechanisms. Nat Metab. 2021;3:166–181. doi: 10.1038/s42255-020-00338-8

15. Liang G, Wang S, Shao J, Jin YJ, Xu L, Yan Y, Günther S, Wang L, Offermanns S. Tenascin-X Mediates Flow-Induced Suppression of EndMT and Atherosclerosis. Circ Res. 2022;130:1647–1659. doi: 10.1161/circresaha.121.320694

16. Mehta V, Pang KL, Rozbesky D, Nather K, Keen A, Lachowski D, Kong Y, Karia D, Ameismeier M, Huang J, et al. The guidance receptor plexin D1 is a mechanosensor in endothelial cells. Nature. 2020;578:290–295. doi: 10.1038/s41586-020-1979-4

17. Dong Y, Wang B, Du M, Zhu B, Cui K, Li K, Yuan K, Cowan DB, Bhattacharjee S, Wong S, et al. Targeting Epsins to Inhibit Fibroblast Growth Factor Signaling While Potentiating Transforming Growth Factor-β Signaling Constrains Endothelial-to-Mesenchymal Transition in Atherosclerosis. Circulation. 2023;147:669–685. doi: 10.1161/circulationaha.122.063075

18. Shelton EL, Yutzey KE. Twistl function in endocardial cushion cell proliferation, migration, and differentiation during heart valve development. Developmental Biology. 2008;317:282–295. doi: 10.1016/j.ydbio.2008.02.037

19. Hunyenyiwa T, Hendee K, Matus K, Kyi P, Mammoto T, Mammoto A. Obesity Inhibits Angiogenesis Through TWIST1-SLIT2 Signaling. Front Cell Dev Biol. 2021;9:693410. doi: 10.3389/fcell.2021.693410

20. Mammoto T, Muyleart M, Konduri GG, Mammoto A. Twist1 in Hypoxia-induced Pulmonary Hypertension through Transforming Growth Factor-β-Smad Signaling. Am J Respir Cell Mol Biol. 2018;58:194–207. doi: 10.1165/rcmb.2016-0323OC

21. Lovisa S, Fletcher-Sananikone E, Sugimoto H, Hensel J, Lahiri S, Hertig A, Taduri G, Lawson E, Dewar R, Revuelta I, et al. Endothelial-to-mesenchymal transition compromises vascular integrity to induce Myc-mediated metabolic reprogramming in kidney fibrosis. Sci Signal. 2020;13. doi: 10.1126/scisignal.aaz2597

22. Kalluri AS, Vellarikkal SK, Edelman ER, Nguyen L, Subramanian A, Ellinor PT, Regev A, Kathiresan S, Gupta RM. Single-Cell Analysis of the Normal Mouse Aorta Reveals Functionally Distinct Endothelial Cell Populations. Circulation. 2019;140:147–163. doi: 10.1161/circulationaha.118.038362

23. Souilhol C, Ayllon BT, Li XY, Diagbouga MR, Zhou ZQ, Canham L, Roddie H, Pirri D, Chambers EV, Dunning MJ, et al. JAG1-NOTCH4 mechanosensing drives atherosclerosis. Science Advances. 2022;8. doi: 10.1126/sciadv.abo7958

24. Blackburn PR, Xu Z, Tumelty KE, Zhao RW, Monis WJ, Harris KG, Gass JM, Cousin MA, Boczek NJ, Mitkov MV, et al. Bi-allelic Alterations in AEBP1 Lead to Defective Collagen Assembly and Connective Tissue Structure Resulting in a Variant of Ehlers-Danlos Syndrome. Am J Hum Genet. 2018;102:696–705. doi: 10.1016/j.ajhg.2018.02.018

25. Welch-Reardon KM, Wu N, Hughes CC. A role for partial endothelial-mesenchymal transitions in angiogenesis? Arteriosclerosis, thrombosis, and vascular biology. 2015;35:303–308. doi: 10.1161/atvbaha.114.303220

26. Fang JS, Hultgren NW, Hughes CCW. Regulation of Partial and Reversible Endothelial-to-Mesenchymal Transition in Angiogenesis. Front Cell Dev Biol. 2021;9:702021. doi: 10.3389/fcell.2021.702021

27. Suzuki T, Carrier EJ, Talati MH, Rathinasabapathy A, Chen X, Nishimura R, Tada Y, Tatsumi K, West J. Isolation and characterization of endothelial-to-mesenchymal transition cells in pulmonary arterial hypertension. Am J Physiol Lung Cell Mol Physiol. 2018;314:L118–l126. doi: 10.1152/ajplung.00296.2017

28. Tombor LS, John D, Glaser SF, Luxán G, Forte E, Furtado M, Rosenthal N, Baumgarten N, Schulz MH, Wittig J, et al. Single cell sequencing reveals endothelial plasticity with transient mesenchymal activation after myocardial infarction. Nat Commun. 2021;12:681. doi: 10.1038/s41467-021-20905-1

29. Kalucka J, de Rooij L, Goveia J, Rohlenova K, Dumas SJ, Meta E, Conchinha NV, Taverna F, Teuwen LA, Veys K, et al. Single-Cell Transcriptome Atlas of Murine Endothelial Cells. Cell. 2020;180:764–779.e720. doi: 10.1016/j.cell.2020.01.015

30. Becker LM, Chen SH, Rodor J, de Rooij L, Baker AH, Carmeliet P. Deciphering endothelial heterogeneity in health and disease at single-cell resolution: progress and perspectives. Cardiovasc Res. 2023;119:6–27. doi: 10.1093/cvr/cvac018

31. Liu JY, Jiang L, Liu JJ, He T, Cui YH, Qian F, Yu PW. AEBP1 promotes epithelial-mesenchymal transition of gastric cancer cells by activating the NF-κB pathway and predicts poor outcome of the patients. Sci Rep. 2018;8:11955. doi: 10.1038/s41598-018-29878-6

32. Fazilaty H, Rago L, Kass Youssef K, Ocaña OH, Garcia-Asencio F, Arcas A, Galceran J, Nieto MA. A gene regulatory network to control EMT programs in development and disease. Nature Communications. 2019;10:5115. doi: 10.1038/s41467-019-13091-8

33. Leroy P, Mostov KE. Slug is required for cell survival during partial epithelial-mesenchymal transition of HGF-induced tubulogenesis. Mol Biol Cell. 2007;18:1943–1952. doi: 10.1091/mbc.e06-09-0823

34. Girard BJ, Knutson TP, Kuker B, McDowell L, Schwertfeger KL, Ostrander JH. Cytoplasmic Localization of Proline, Glutamic Acid, Leucine-rich Protein 1 (PELP1) Induces Breast Epithelial Cell Migration through Up-regulation of Inhibitor of κB Kinase ɛ and Inflammatory Cross-talk with Macrophages. J Biol Chem. 2017;292:339–350. doi: 10.1074/jbc.M116.739847

35. Roy SS, Gonugunta VK, Bandyopadhyay A, Rao MK, Goodall GJ, Sun LZ, Tekmal RR, Vadlamudi RK. Significance of PELP1/HDAC2/miR-200 regulatory network in EMT and metastasis of breast cancer. Oncogene. 2014;33:3707–3716. doi: 10.1038/onc.2013.332

36. Holloway RW, Bogachev O, Bharadwaj AG, McCluskey GD, Majdalawieh AF, Zhang L, Ro HS. Stromal adipocyte enhancer-binding protein (AEBP1) promotes mammary epithelial cell hyperplasia via proinflammatory and hedgehog signaling. J Biol Chem. 2012;287:39171–39181. doi: 10.1074/jbc.M112.404293

37. Corano Scheri K, Lavine JA, Tedeschi T, Thomson BR, Fawzi AA. Single-cell transcriptomics analysis of proliferative diabetic retinopathy fibrovascular membranes reveals AEBP1 as fibrogenesis modulator. JCI Insight. 2023;8. doi: 10.1172/jci.insight.172062

38. Kattih B, Boeckling F, Shumliakivska M, Tombor L, Rasper T, Schmitz K, Hoffmann J, Nicin L, Abplanalp WT, Carstens DC, et al. Single-nuclear transcriptome profiling identifies persistent fibroblast activation in hypertrophic and failing human hearts of patients with longstanding disease. Cardiovasc Res. 2023;119:2550–2562. doi: 10.1093/cvr/cvad140

39. Wang D, Rabhi N, Yet SF, Farmer SR, Layne MD. Aortic carboxypeptidase-like protein regulates vascular adventitial progenitor and fibroblast differentiation through myocardin related transcription factor A. Sci Rep. 2021;11:3948. doi: 10.1038/s41598-021-82941-7

40. Tumelty KE, Smith BD, Nugent MA, Layne MD. Aortic carboxypeptidase-like protein (ACLP) enhances lung myofibroblast differentiation through transforming growth factor β receptor-dependent and-independent pathways. J Biol Chem. 2014;289:2526–2536. doi: 10.1074/jbc.M113.502617

41. Steffensen LB, Rasmussen LM. A role for collagen type IV in cardiovascular disease? Am J Physiol Heart Circ Physiol. 2018;315:H610–h625. doi: 10.1152/ajpheart.00070.2018

42. McNeilly S, Thomson CR, Gonzalez-Trueba L, Sin YY, Granata A, Hamilton G, Lee M, Boland E, McClure JD, Lumbreras-Perales C, et al. Collagen IV deficiency causes hypertrophic remodeling and endothelium-dependent hyperpolarization in small vessel disease with intracerebral hemorrhage. EBioMedicine. 2024;107:105315. doi: 10.1016/j.ebiom.2024.105315

43. Steffensen LB, Stubbe J, Lindholt JS, Beck HC, Overgaard M, Bloksgaard M, Genovese F, Holm Nielsen S, Tha MLT, Bang-Moeller SK, et al. Basement membrane collagen IV deficiency promotes abdominal aortic aneurysm formation. Sci Rep. 2021;11:12903. doi: 10.1038/s41598-021-92303-y

44. Jin R, Shen J, Zhang T, Liu Q, Liao C, Ma H, Li S, Yu Z. The highly expressed COL4A1 genes contributes to the proliferation and migration of the invasive ductal carcinomas. Oncotarget. 2017;8:58172–58183. doi: 10.18632/oncotarget.17345

45. Zhang H, Wang Y, Ding H. COL4A1, negatively regulated by XPD and miR-29a-3p, promotes cell proliferation, migration, invasion and epithelial-mesenchymal transition in liver cancer cells. Clin Transl Oncol. 2021;23:2078–2089. doi: 10.1007/s12094-021-02611-y

46. Large-scale gene-centric analysis identifies novel variants for coronary artery disease. PLoS Genet. 2011;7:e1002260. doi: 10.1371/journal.pgen.1002260

47. Malik R, Rannikmäe K, Traylor M, Georgakis MK, Sargurupremraj M, Markus HS, Hopewell JC, Debette S, Sudlow CLM, Dichgans M. Genome-wide meta-analysis identifies 3 novel loci associated with stroke. Ann Neurol. 2018;84:934–939. doi: 10.1002/ana.25369

48. Wentzel JJ, Bos D, White SJ, van der Heiden K, Kavousi M, Evans PC. Sex-related differences in coronary and carotid vessel geometry, plaque composition and shear stress obtained from imaging. Atherosclerosis. 2024;395:117616. doi: 10.1016/j.atherosclerosis.2024.117616

49. Sakkers TR, Mokry M, Civelek M, Erdmann J, Pasterkamp G, Diez Benavente E, den Ruijter HM. Sex differences in the genetic and molecular mechanisms of coronary artery disease. Atherosclerosis. 2023;384:117279. doi: 10.1016/j.atherosclerosis.2023.117279

50. Shin J, Hong J, Edwards-Glenn J, Krukovets I, Tkachenko S, Adelus ML, Romanoski CE, Rajagopalan S, Podrez E, Byzova TV, et al. Unraveling the Role of Sex in Endothelial Cell Dysfunction: Evidence From Lineage Tracing Mice and Cultured Cells. *Arteriosclerosis*, Thrombosis, and Vascular Biology. 2024;44:238–253. doi: 10.1161/ATVBAHA.123.319833

51. Tardajos Ayllon B, Bowden N, Souilhol C, Darwish H, Tian S, Duckworth C, Pritchard DM, Xu S, Sayers J, Francis S, et al. Endothelial c-REL orchestrates atherosclerosis at regions of disturbed flow through crosstalk with TXNIP-p38 and non-canonical NF-κB pathways. Cardiovasc Res. 2025. doi: 10.1093/cvr/cvaf024

52. Sun J, Singh P, Shami A, Kluza E, Pan M, Djordjevic D, Michaelsen NB, Kennbäck C, van der Wel NN, Orho-Melander M, et al. Spatial Transcriptional Mapping Reveals Site-Specific Pathways Underlying Human Atherosclerotic Plaque Rupture. J Am Coll Cardiol. 2023;81:2213–2227. doi: 10.1016/j.jacc.2023.04.008

53. Singh P, Sun J, Cavalera M, Al-Sharify D, Matthes F, Barghouth M, Tengryd C, Dunér P, Persson A, Sundius L, et al. Dysregulation of MMP2-dependent TGF-ß2 activation impairs fibrous cap formation in type 2 diabetes-associated atherosclerosis. Nat Commun. 2024;15:10464. doi: 10.1038/s41467-024-50753-8

54. Robinson MD, McCarthy DJ, Smyth GK. edgeR: a Bioconductor package for differential expression analysis of digital gene expression data. Bioinformatics. 2010;26:139–140. doi: 10.1093/bioinformatics/btp616

55. Johnson WE, Li C, Rabinovic A. Adjusting batch effects in microarray expression data using empirical Bayes methods. Biostatistics. 2007;8:118–127. doi: 10.1093/biostatistics/kxj037

