## Supplementl information for "TWIST1 drives endothelial-to-mesenchymal-transition to stabilize atherosclerotic plaques"

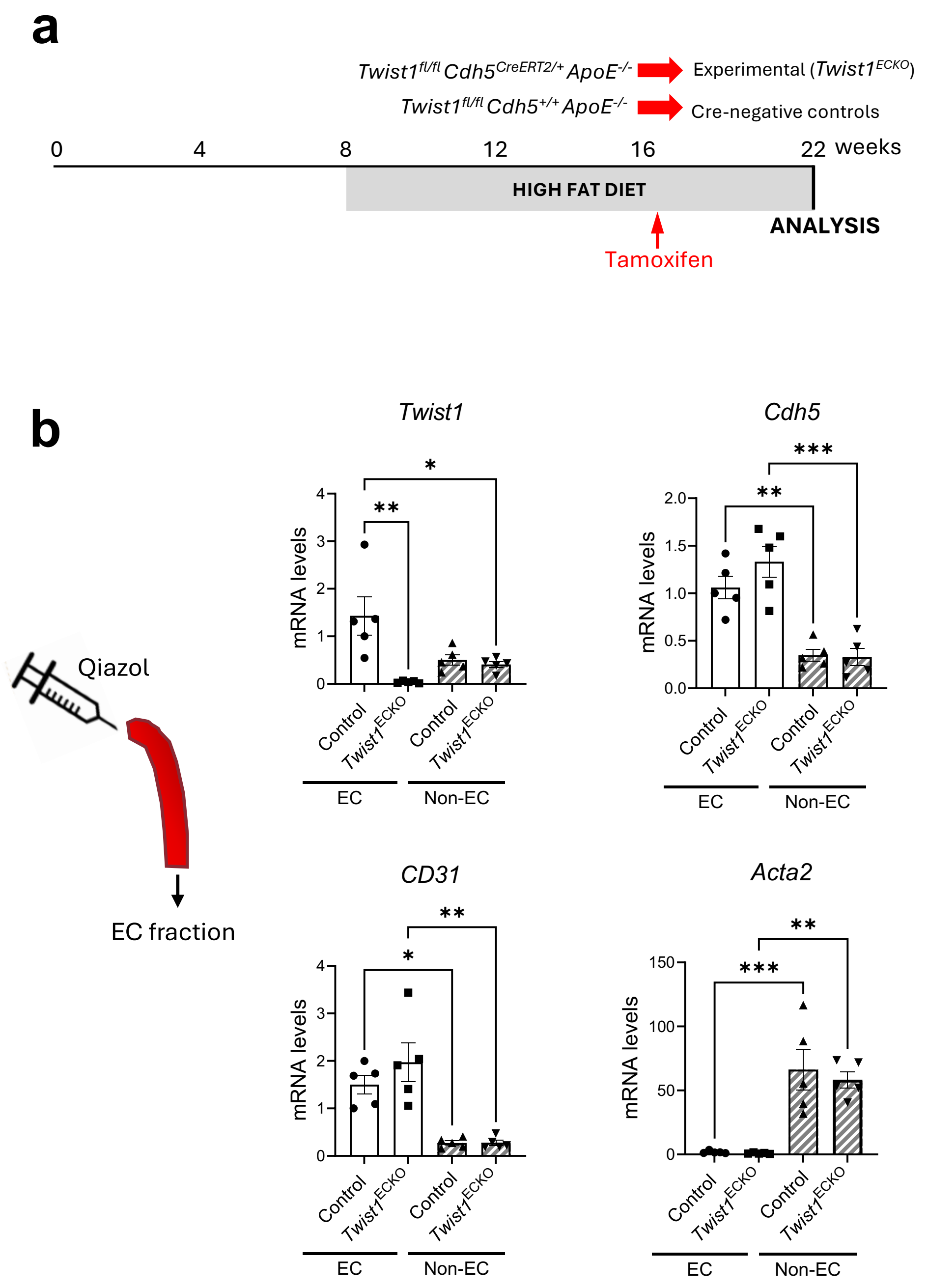


**Figure S1. Validation of *Twist1* knock-down in mouse aorta.** (A) Timeline of *Twist1* deletion in a model of atherosclerotic progression. *Twist1*$^{ECKO}$ (*Twist1*$^{fl/fl}$ *Cdh5*$^{CreERT2/+}$ *ApoE*$^{-/-}$) and control mice (*Twist1*$^{fl/fl}$ *Cdh5*$^{+/+}$ *ApoE*$^{-/-}$) aged 8 weeks were fed a Western diet for 8 weeks to induce atherosclerotic lesions. At that point, tamoxifen was administered for 5 consecutive days to induce *Twist1* deletion and a Western diet was provided for an additional 6 weeks (totalling 14 weeks of Western diet). (B) Validation of *Twist1* deletion in the endothelium. RNA was extracted from EC fractions (by flushing of Qiazol) and from residual medial/adventitial tissue (Non-EC) of aortas isolated from *Twist1*$^{ECKO}$ and control mice. Expression levels of *Twist1, Cdh5, CD31* and *Acta2* were quantified by qRT-PCR (n=5). Mean values are shown +/- standard errors. Differences between means were analysed using a 2-way ANOVA.

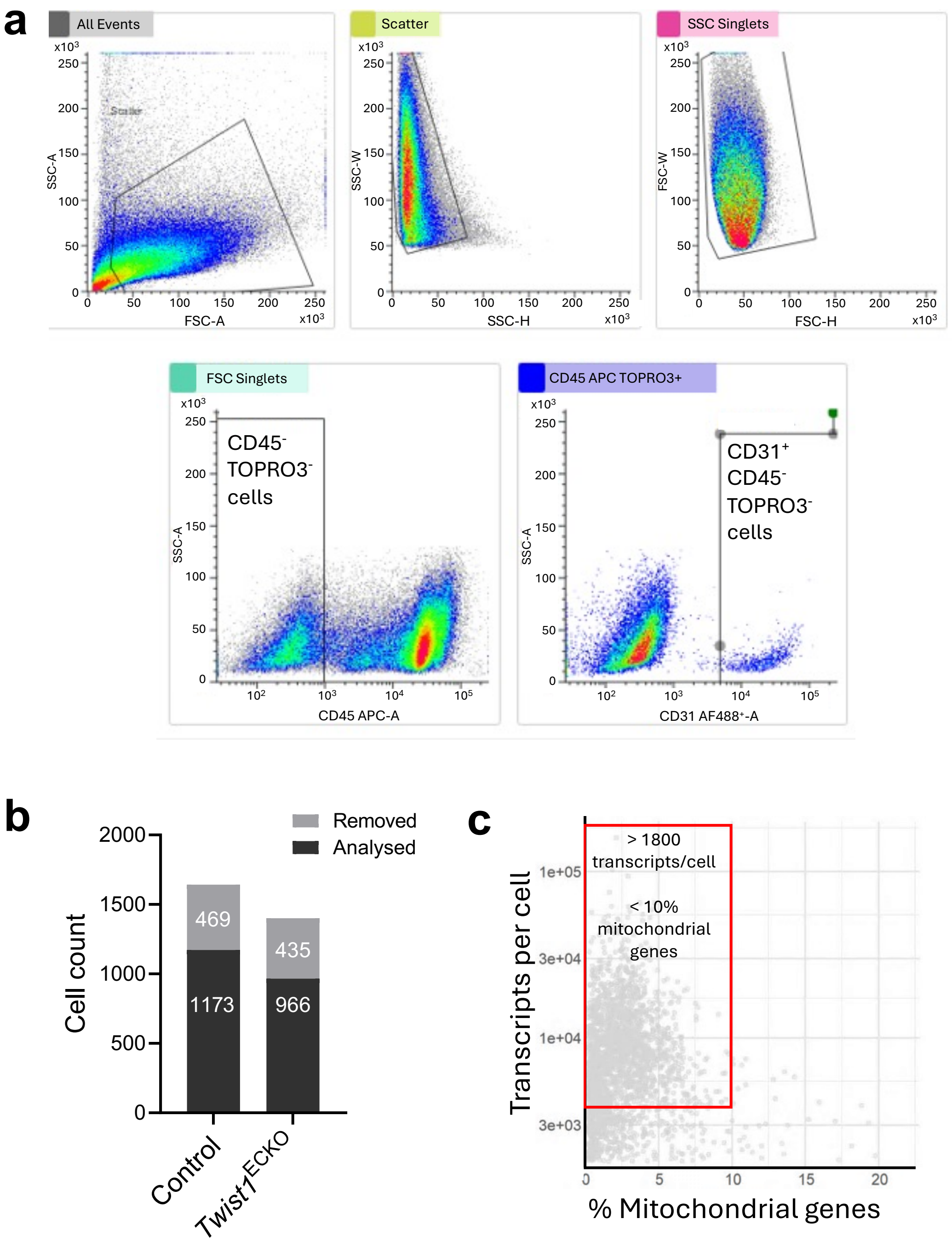


**Figure S2. scRNAseq analysis sorting strategy and quality control.** Aortas from *Twist1* $EC^{KO}$ and control mice after 14 weeks of Western diet were analysed by FACS of CD31$^+$ CD45$^-$ cells coupled to scRNA-seq. (A) Representative flow cytometry workflow and sorting strategy to isolate of CD31$^+$/CD45$^-$/TOPRO3$^-$ cells. Single cells were identified through side scatter and forward scatter and the CD45-/TOPRO3- population was subsequently selected. From the CD45$^-$/TOPRO3$^-$ population, CD31$^+$ cells were sorted into a 384-well plate for scRNAseq. (B) Bar graph showing analysed cells by genotype after quality control (QC) analysis. Cells that were removed during QC analysis are represented in light grey, whereas remaining cells after QC filtering are represented in black. (C) Scatter plot showing transcripts per cell vs % Mitochondrial genes. Cells inside the red box, with <10% mitochondrial content and >1800 transcripts/cell, were selected and used for subsequent scRNAseq.

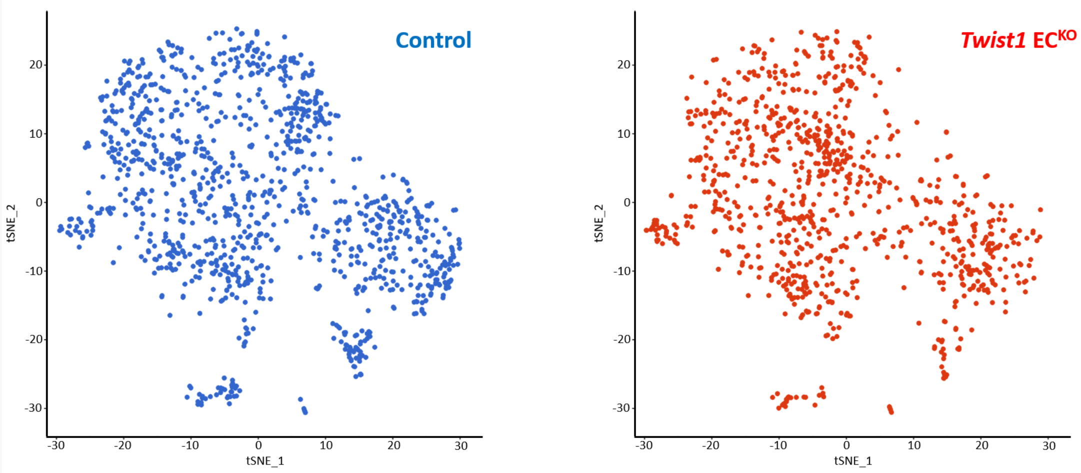


**Figure S3. EC from *Twist1$^{ECKO}$* and control mice exhibit different clustering patterns by scRNAseq.** Aortas from *Twist1$^{ECKO}$* and control mice after 14 weeks of Western diet were analysed by FACS of CD31$^{+}$ CD45$^{-}$ cells coupled to scRNA-seq. t-SNE maps showing the distribution of aortic CD31$^{+}$ CD45$^{-}$ cells from control (left) and *Twist1$^{ECKO}$* (right) mice.

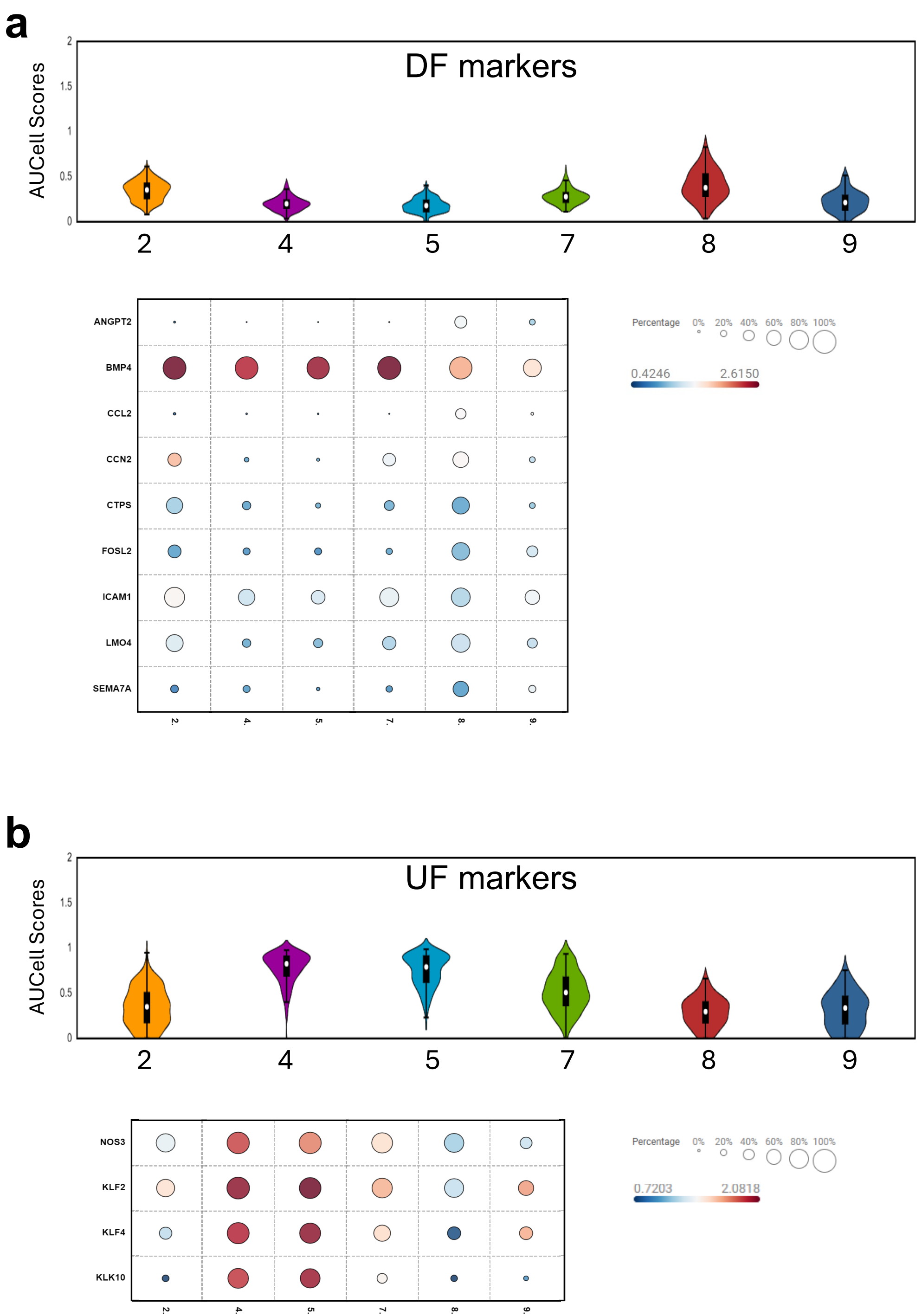


**Figure S4. Expression of uniform flow and disturbed flow markers in selected scRNAseq clusters**. After 14 weeks of Western diet, $CD31^{+}$ $CD45^{-}$ cells from aortas from *Twist1* $EC^{KO}$ and control mice were processed for scRNA-seq. (A) DF markers were measured in each cell. At the top, DF markers are presented as a violin plot as an average in selected clusters. At the bottom, DF markers are presented individually as a bubble plot in selected clusters. (B) Uniform flow (UF) markers were measured in each cell. At the top, UF markers are presented as a violin plot as an average in selected clusters. At the bottom, UF markers are presented individually as a bubble plot in selected clusters. Clusters 2, 7, 8 and 9 (largely composed of ECs derived from control mice) and 4 and 5 (mainly composed of ECs derived from *Twist1*$^{ECKO}$ mice were compared.

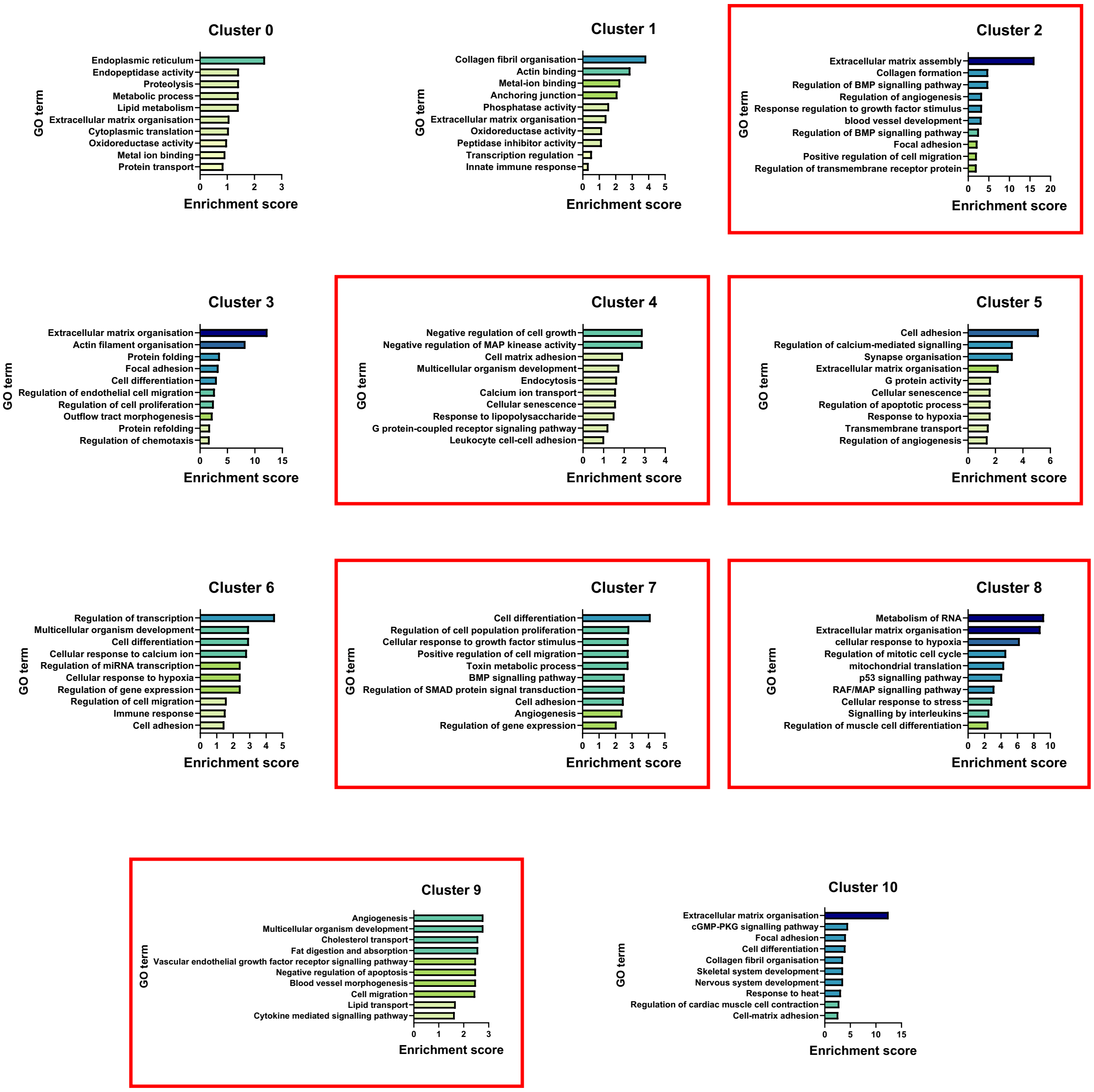


**Figure S5. Functional annotation of scRNAseq.** scRNA-seq analysis of CD31$^+$ CD45$^-$ cells from *Twist1*$^{ECKO}$ and control mice revealed 11 distinct clusters. The most highly enriched GO pathways for each cluster, as well as their enrichment scores, are shown. Clusters 2, 7, 8 and 9 (largely composed of ECs derived from control mice) and 4 and 5 (mainly composed of ECs derived from *Twist1*$^{ECKO}$ mice) are highlighted in red.

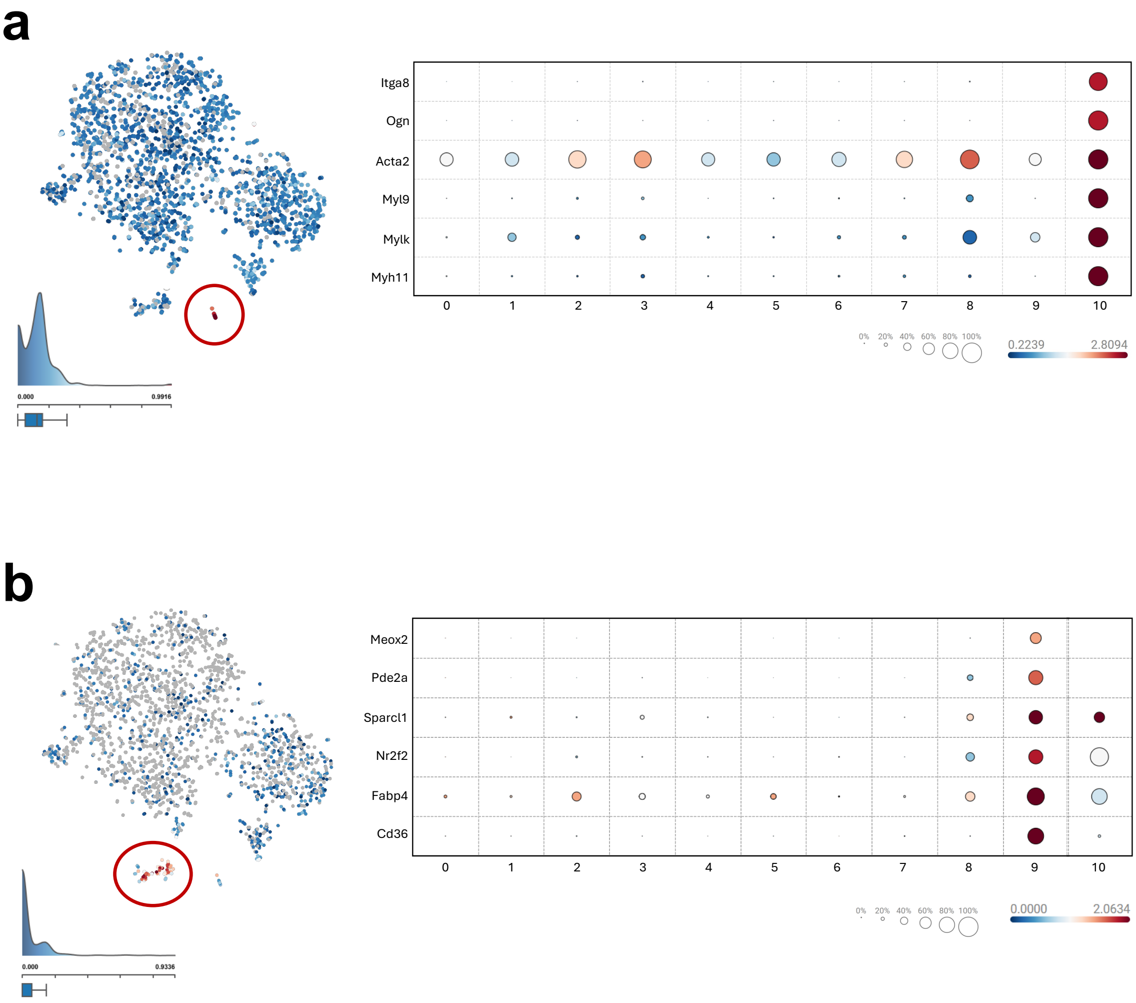


**Figure S6. Expression of enriched markers in scRNAseq clusters 9 and 11.** scRNA-seq analysis of CD31$^{+}$ CD45$^{-}$ cells from *Twist1$^{ECKO}$* and control mice revealed 11 distinct clusters. (A) t-SNE map and bubble map indicating the expression of the most highly enriched markers in cluster 9. Cluster 9 is circled in red on the t-SNE map. (B) t-SNE map and bubble map indicating the expression of the most highly enriched markers in cluster 10. Cluster 10 is circled in red on the t-SNE map.

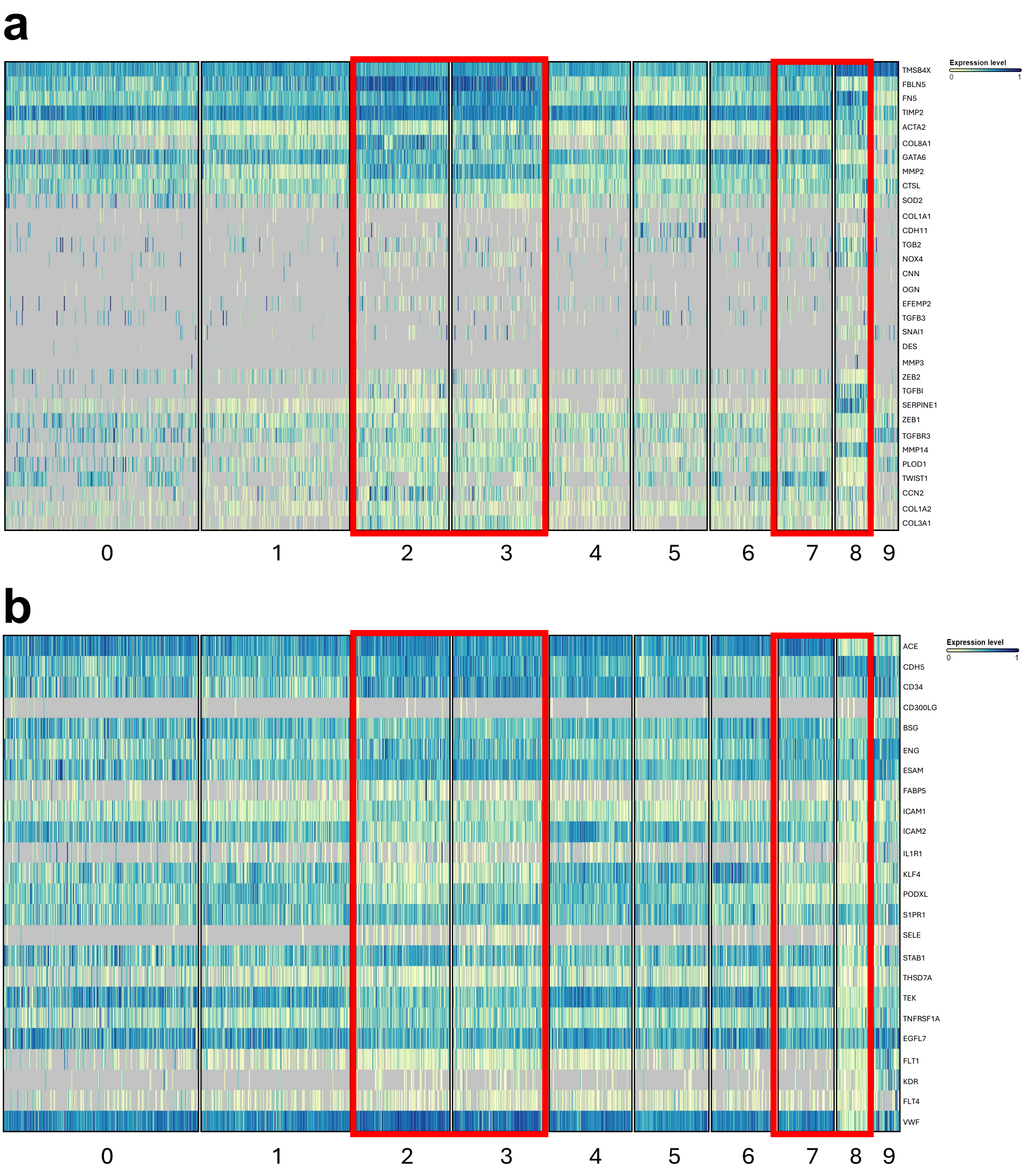


**Figure S7**. **Expression of markers of EndMT-like cells and ECs markers in scRNAseq clusters.** Aortas from *Twist1*$^{ECKO}$ and control mice after 14 weeks of Western diet were analysed by FACS of CD31$^{+}$ CD45$^{-}$ cells coupled to scRNA-seq. (A) Heatmap showing expression of markers of EndMT in clusters 0-9 (EC clusters). (B) Markers of ECs in clusters 0-9. Clusters 2, 3, 7 and 8, which show enriched expression of EndMT markers, are highlighted in red.

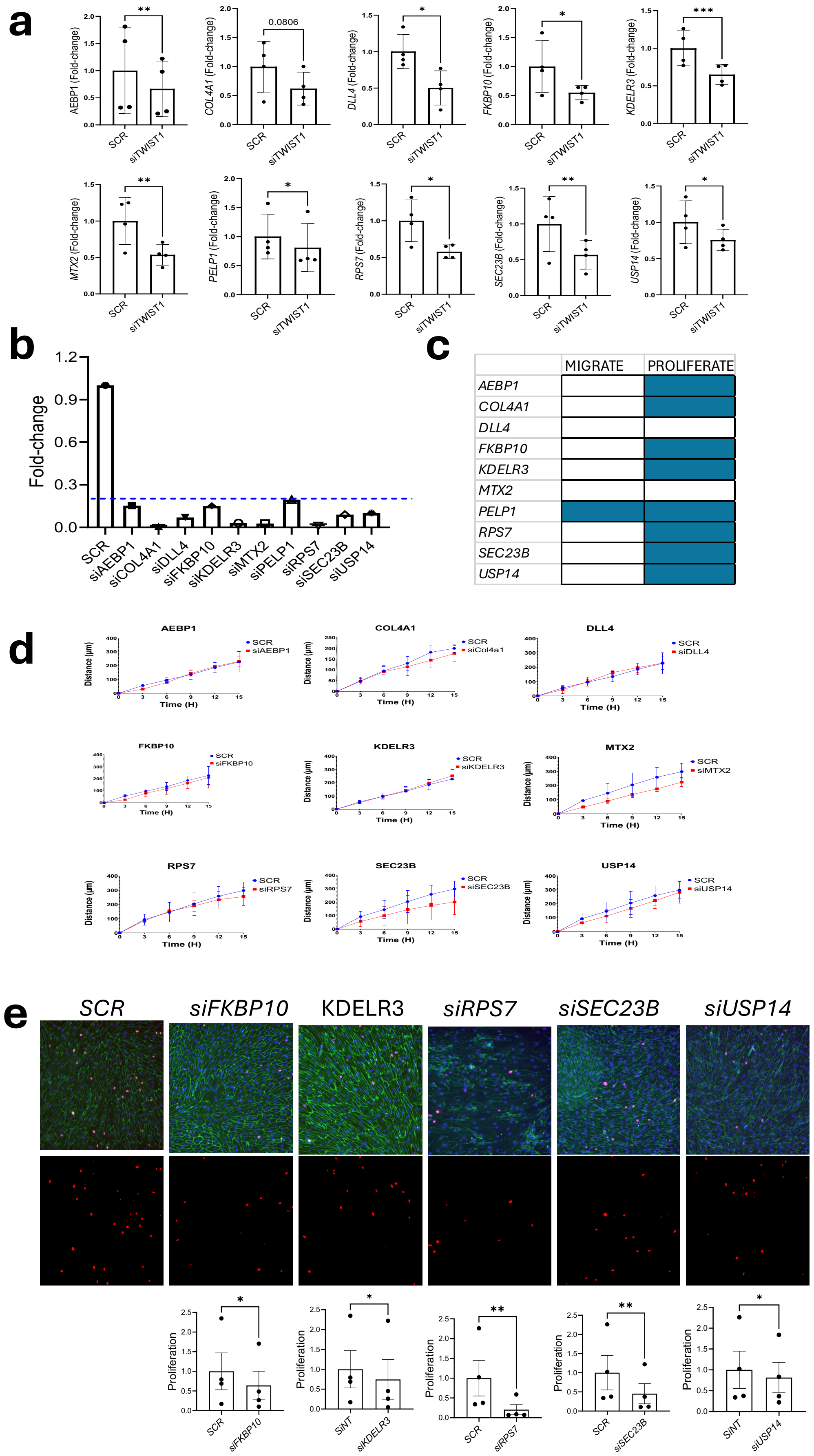
a
AEBP1 (Fold-change)
COL4A1 (Fold-change)
DLL4 (Fold-change)
FKBP10 (Fold-change)
KDELR3 (Fold-change)
MTX2 (Fold-change)
PELP1 (Fold-change)
RPS7 (Fold-change)
SEC23B (Fold-change)
USP14 (Fold-change)
SCR
siTWIST1
0.0806
b
Fold-change
SCR
siAEBP1
siCOL4A1
siDLL4
siFKBP10
siKDELR3
siMTX2
siPELP1
siRPS7
siSEC23B
siUSP14
c
MIGRATE
PROLIFERATE
AEBP1
COL4A1
DLL4
FKBP10
KDELR3
MTX2
PELP1
RPS7
SEC23B
USP14
d
Distance (μm)
Time (H)
siAEBP1
siCol4a1
siDLL4
siFKBP10
siKDELR3
siMTX2
siRPS7
siSEC23B
siUSP14
e
SCR
siFKBP10
KDELR3
siRPS7
siSEC23B
siUSP14
Proliferation
SiNT
siKDELR3

**Figure S8. TWIST1 regulated genes are involved in cell migration and proliferation.** (A) qPCR validation of RNA-seq data. Gene expression levels of *AEBP1*, *COL4A1*, *DLL4*, *FKBP10*, *KDELR3*, *MTX2*, *PELP1*, *RPS7*, *USP14*, and *SEC23B* in *TWIST1*-silenced HAECs compared to control (*SCR*) cells (n=4). (B) qPCR confirmation of siRNA-mediated silencing of these genes. (C) Summary table categorizing genes based on their involvement in proliferation and/or migration. (D) Cell migration was assessed using a scratch wound assay in HAEC monolayers transfected with siRNA targeting *AEBP1*, *COL4A1*, *DLL4*, *FKBP10*, *KDELR3*, *MTX2*, *RPS7*, *USP14*, and *SEC23B,* or a non-targeting control (SCR) after exposure to DF for 72h (orbital shaker). Quantification of migration distance from the initial wound (T0) at multiple time points is shown. (E) Ki67 immunofluorescence staining (red) was performed to assess proliferation after siRNA-mediated silencing of *FKBP10-*, RPS7 and *SEC23* in HAECs exposed to DF for 72h (orbital shaker). Merged images show DAPI (blue) and CDH5 (green) (scale bar= 100 µm). Ki67-positive cells were quantified as a proportion of total nuclei and values are shown as relative fold change compared to *SCR* (n=4). Mean values are shown +/- standard errors. Differences between means were analysed using a ratio paired t-test.

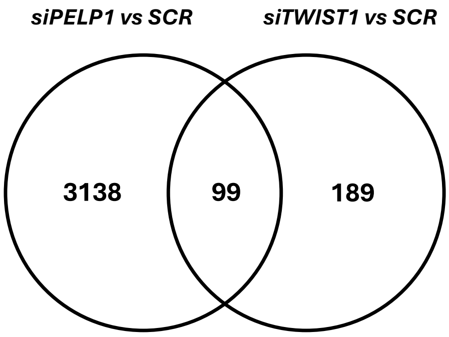


**99 genes co-regulated by PELP1 and TWIST1**

| | | | | | | |
|---|---|---|---|---|---|---|
| ADPRH | CMTM3 | FRS2 | LOC64413 | OAS2 | RNF152 | VPS54 |
| AEBP1 | CNOT6 | **GRB10** | 5 | P3H1 | RPS25 | WASHC4 |
| **ANXA3** | COL4A1 | GSDMC | LRRC8A | **PELP1** | **SEMA3A** | WDFY4 |
| ATP9A | CREBRF | HADHA | MAGED4B | PHYH | SLC12A2 | ZNF24 |
| BCL7A | **CRKL** | **HES1** | MAP1A | PLA2G12A | SLC35D1 | |
| BDH1 | **CSF1** | HLA-DRB1 | MAPK6 | PLIN3 | SNCA | |
| C17orf58 | CYB561A3 | HMGA2 | MEDAG | PNRC2 | **SPOCK1** | |
| C5orf15 | DANCR | HSPA4L | MFAP3 | **PODN** | SPR | |
| CASP4 | **DPP4** | **ICAM1** | MFNG | PPP1R18 | STMN3 | |
| CASP7 | ECE1 | IFI27 | MRC2 | **PTK7** | STRADB | |
| CCPG1 | EEF2 | IFITM3 | MSL2 | RABAC1 | SV2A | |
| **CEACAM1** | EIF4EBP2 | **IGSF8** | MTX2 | RCC1L | SYT11 | |
| CELF2 | **EMILIN1** | **ITGB4** | NACC2 | RCN3 | TMEM184B | |
| CHST1 | ERC1 | KDELR3 | NCSTN | RENBP | TOB1 | |
| CLCN3 | FKBP10 | LAMB2 | NUCB1 | RFX2 | TOR1A | |
| CLCN4 | FRMD4B | LINC00520 | OAS1 | **RHOU** | VPS4B | |

**Figure S9. TWIST1 and PELP1 co-regulate genes that promote EC migration.** RNA-seq analysis was performed after siRNA-mediated silencing of *TWIST1* or *PELP1* in HAECs exposed to DF for 72h (orbital shaker). Venn diagram illustrating genes co-regulated by *TWIST1 and PELP1* and those regulated exclusively by these molecules. On the right, genes co-regulated by *TWIST1* and *PELP1* are listed with migration-associated genes highlighted in red.

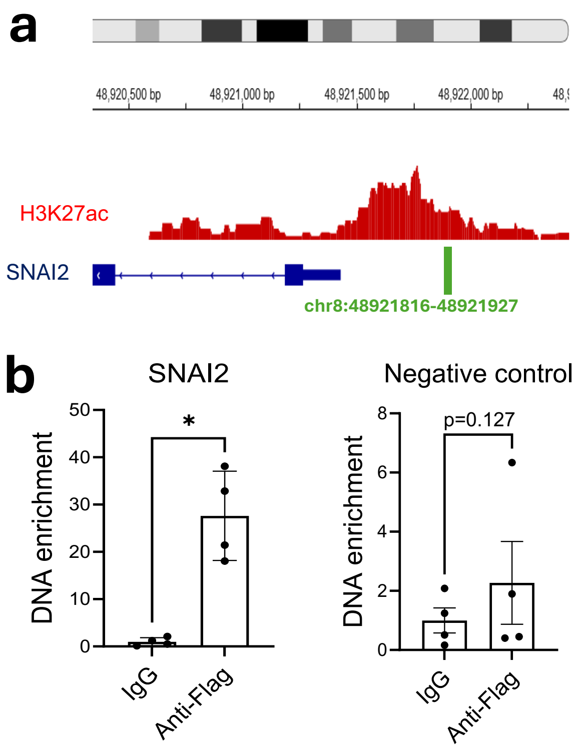


**Figure S10. ChIP-qPCR analysis of TWIST1 in HAECs under disturbed flow.** HAECs were infected with lentivirus expressing TWIST1-FLAG and exposed to DF using an orbital shaker for 72h. TWIST1 binding at the *SNAI2* locus was assessed using an anti-FLAG antibody and using IgG as a control. (A) Schematic representation of the *SNAI2* gene loci, with TWIST1 binding sites highlighted in green. (B) ChIP-qPCR revealed enrichment of *SNAI2* regulatory region DNA to the IgG control (left), whereas irrelevant control sequences were not enriched (n=4). Mean values are shown +/- standard errors. Differences between means were analysed using a ratio paired t-test.

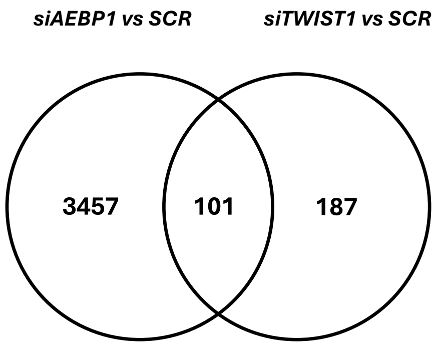


**101 genes co-regulated by AEBP1 and TWIST1**

| | | | | | |
|---|---|---|---|---|---|
| ADPRH | CLCN4 | EXT2 | **LAMB2** | **PODN** | STMN3 |
| AEBP1 | CLN8 | FAM168B | LTBP1 | PREX1 | SUMO3 |
| ANXA3 | CNOT6 | FILIP1L | LZIC | PRX | SUPT6H |
| **APP** | **COL18A1** | FRMD4B | MAGED4B | PTGS1 | **SUZ12** |
| ARSB | COL4A1 | **FRS2** | MAPK6 | PTK7 | SV2A |
| ATP9A | **CRKL** | GPX7 | MEDAG | RALBP1 | SYT11 |
| B3GNT9 | **CSF1** | GSTM4 | MFNG | **RCN3** | TMCC3 |
| **BCL7A** | CSNK1G3 | **HES1** | MRC2 | RENBP | TMEM255B |
| BDH1 | DANCR | **HMGA2** | MTX2 | RPL10A | TSPAN17 |
| C5orf15 | **DLL4** | HSPA4L | **NACC2** | RPL28 | UNC119B |
| C6orf120 | DPP4 | ICAM1 | **NCSTN** | **SDE2** | WASHC4 |
| CASP4 | EBPL | IGF2BP2 | OAS2 | SEC23B | **WEE1** |
| CASP7 | EIF4EBP2 | IGSF8 | **P3H3** | SEMA3A | WRNIP1 |
| **CDK19** | **EMILIN1** | ITGB4 | PAPLN | SLC12A2 | ZNF532 |
| CEACAM1 | **ENSA** | KDELR3 | PCSK5 | SLC16A3 | ZNF678 |
| CHST1 | ERC1 | KDM6B | **PDCD1LG2** | SPOCK1 | **ZNF703** |
| CHST12 | ERMP1 | KLF12 | PLIN3 | SPRY4 | |

**Figure S11. TWIST1 and AEBP1 co-regulate genes that promote EC proliferation.** RNA-seq analysis was performed after siRNA-mediated silencing of *TWIST1* or *AEBP1* in HAECs exposed to DF for 72h (orbital shaker). Venn diagram illustrating genes co-regulated by *TWIST1 and AEBP1* and those regulated exclusively by these molecules. On the right, genes co-regulated by *TWIST1* and *AEBP1* are listed with proliferation-associated genes highlighted in blue.

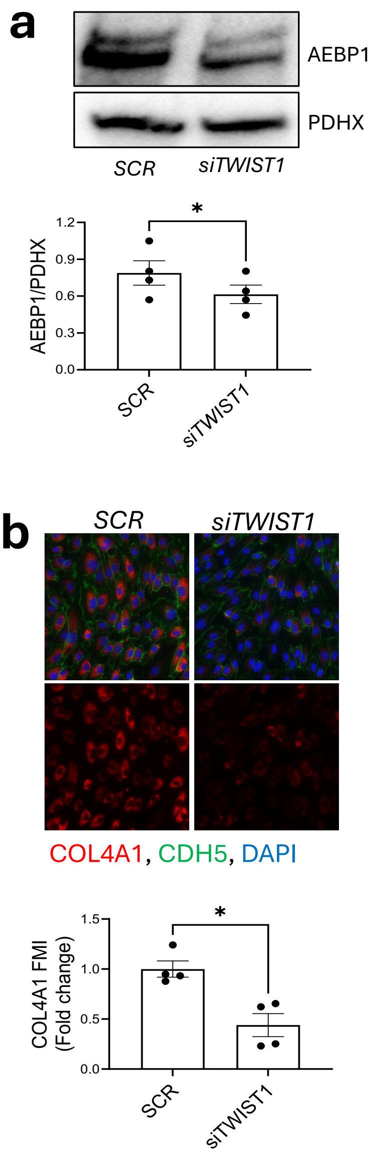


**Figure S12. TWIST1 silencing reduces AEBP1 and COL4A1 expression in HAECs under DF.** (A) Western blot analysis of AEBP1 expression in *SCR* vs *TWIST1*-silenced HAECs after 72h of DF (Ibidi system), normalized to PDHX (n = 4). (B) Immunofluorescence staining of COL4A1 (red) in HAECs after 72h of DF (Ibidi system), with merged images showing DAPI (nuclear stain, blue) and CDH5 (green) (scale bar= 100µm) (n=4). Mean values are shown +/- standard errors. Differences between means were analysed using a ratio paired t-test.

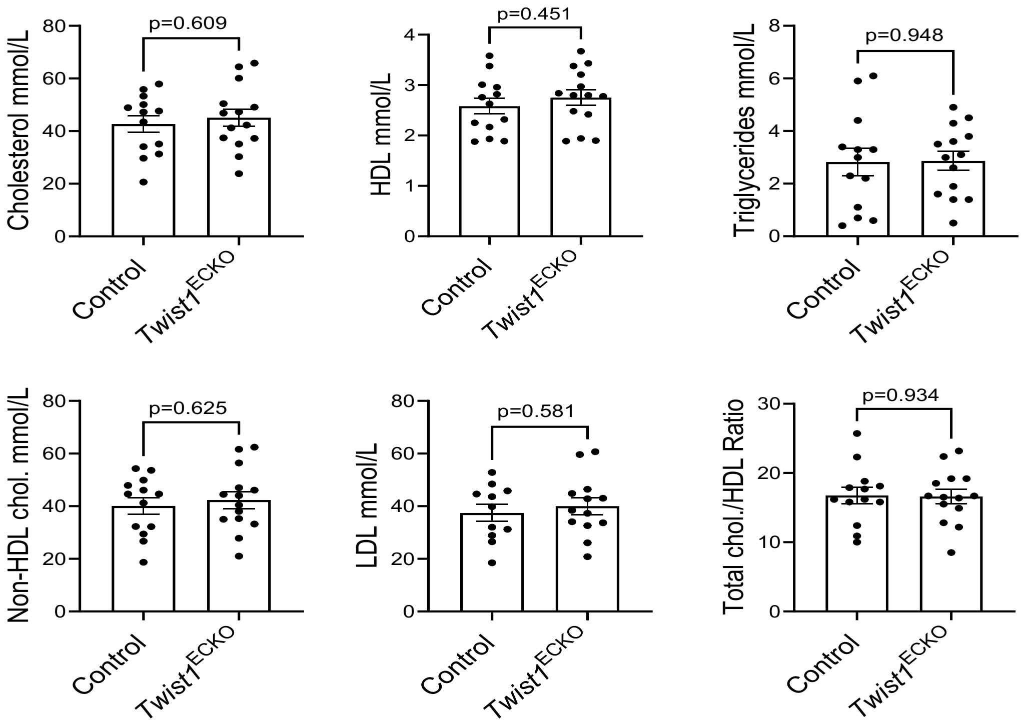


**Figure S13. *Twist1* does not regulate levels of plasma lipoproteins.** *Twist1*$^{ECKO}$ and control mice aged 8 weeks were fed a Western diet for 8 weeks to induce atherosclerotic lesions. Tamoxifen was administered for 5 consecutive days and a Western diet was provided for an additional 6 weeks. Total plasma cholesterol, HDL cholesterol, triglycerides, non-HDL cholesterol, LDL cholesterol and total cholesterol/HDL ratio were measured in *Twist1*$^{ECKO}$ (n=13) and control (n=11) mice. Mean levels +/- standard errors are shown. Differences between means were analysed using an unpaired t-test.

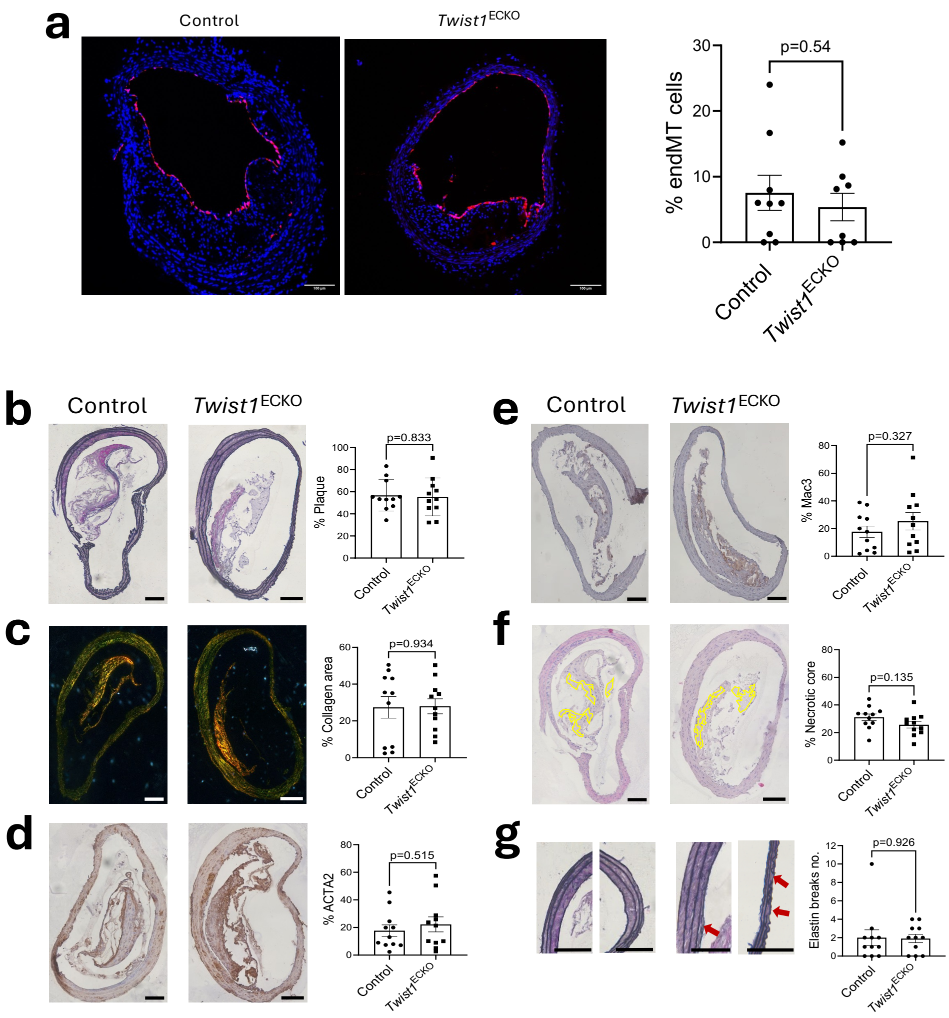


**Figure S14. *Twist1* does not control EndMT, plaque growth or plaque composition in female mice.** (A) Female *Twist1*$^{ECKO}$ (*Twist1*$^{fl/fl}$ *Cdh5*$^{CreERT2/+}$ *ApoE*$^{-/-}$ *Rosa26*$^{TdTomato/TdTomato}$) and control mice (*Twist1*$^{fl/fl}$ *Cdh5*$^{+/+}$ *ApoE*$^{-/-}$ *Rosa26*$^{TdTomato/TdTomato}$) aged 8 weeks were fed a Western diet for 8 weeks to induce atherosclerotic lesions. Tamoxifen was then administered for 5 consecutive days to induce *Twist1* deletion and TdTomato expression in ECs and a Western diet was provided for an additional 6 weeks (totalling 14 weeks of Western diet). The percentage of Rosa26TdTomato$^{+}$ cells that have undergone EndMT was quantified in frozen brachiocephalic sections from *Twist1*$^{ECKO}$ (n=8) and control (n=9) mice. Rosa26TdTomato$^{+}$ cells are shown in red and nuclei are counterstained with DAPI (blue). Representative images are shown (Scale bar=100 μm). (B-F) Female *Twist1*$^{ECKO}$ (*Twist1*$^{fl/fl}$ *Cdh5*$^{CreERT2/+}$ *ApoE*$^{-/-}$) and control mice (*Twist1*$^{fl/fl}$ *Cdh5*$^{+/+}$ *ApoE*$^{-/-}$) aged 8 weeks were fed a Western diet for 8 weeks to induce atherosclerotic lesions. Tamoxifen was then administered for 5 consecutive days to induce *Twist1* deletion in ECs and a Western diet was provided for an additional 6 weeks (totalling 14 weeks of Western diet). Paraffin-embedded sections of brachiocephalic arteries were stained with (B) Miller's elastin stain to quantify plaque burden (C) Picrosirius Red (visualised under polarised light) to quantify collagen content (D) antibodies against ACTA2 to quantify vSMCs content (E) antibodies against MAC3 to quantify macrophage content and (F) Hematoxylin and Eosin (H&E) to quantify necrotic core content (highlighted in yellow) in *Twist1*$^{ECKO}$ (n=11) and control (n=11) mice. (G) Magnified view of regions shown in (B) and quantification of elastin breaks number in brachiocephalic arteries from *Twist1*$^{ECKO}$ (n=11) and control (n=11) mice. The red arrows indicate elastin breaks. Representative images are shown (Scale bar=100 μm). Mean values are shown +/- standard errors. Differences between means were analysed using an unpaired t-test.

### Table S1 PCR Primers for Genotyping

| Primer name | Primer sequence |
|---|---|
| ***Twist1*** Forward | CTTCTCCGTCTGGAGGATGG |
| ***Twist1*** Reverse | GATGGCGTTTTGGGCACAAGG |
| ***Cdh5-Cre*** Forward | TCGATGCAACGAGTGATGAG |
| ***Cdh5-Cre*** Reverse | AGTGCGTTCGAACGCTAGAG |
| ***ApoE*** Null Forward | GCCTAGCCGAGGGAGAGCCG |
| ***ApoE*** WT Forward | TGTGACTTGGGAGCTCTGCAGC |
| ***ApoE*** Reverse | GCCGCCCCGACTGCATCT |
| ***Rosa26TdTomato*** WT Forward | AAGGGAGCTGCAGTGGAGT |
| ***Rosa26TdTomato*** WT Reverse | CCGAAAATCTGTGGGAAGTC |
| ***Rosa26TdTomato*** Mutant Forward | CTGTTCCTGTACGGCATGG |
| ***Rosa26TdTomato*** Mutant Reverse | GGCATTAAAGCAGCGTATCC |

### Table S2 PCR Primers for qRT-PCR

| Organism | Gene | Forward primer | Reverse primer | Purpose |
|---|---|---|---|---|
| *Mus musculus* | *Acta2* | CATCATGCGTCTGGACTTGG | AATCTCACGCTCGGCAGTAG | qRT-PCR |
| *Mus musculus* | *Cdh5* | TCTTGCCAGCAAACTCTCCT | TTGGAATCAAATGCACATCG | qRT-PCR |
| *Mus musculus* | *Cd31* | CGGTGTTCAGCGAGATCC | ACTCGACAGGATGGAAATCAC | qRT-PCR |
| *Mus musculus* | *Hprt* | AGTCCCAGCGTCGTGATTAG | TCTCGAGCAAGTCTTTCAGTCC | qRT-PCR |
| *Mus musculus* | *Twist1* | ACCTAGATGTCATTGTTTCCAGA | CCACGCCCTGATTCTTGTG | qRT-PCR |
| *Homo sapiens* | *AEBP1* | TGAGCGCCAGACAGACGAA | CCTTTCGGGGCTCTTTGTG | qRT-PCR |
| *Homo sapiens* | *COL4A1* | TGCGGCTCAAAGGTGACAA | AATCCTACAGAACCCGGCGA | qRT-PCR |
| *Homo sapiens* | *DLL4* | TCCAACTGCCCTTCAATTTC | ACTGCAGATGACCCGGTAAG | qRT-PCR |
| *Homo sapiens* | *FKBP10* | CATGGGCATGTGTGTCAACG | GAATGAGCCCCGCCAGG | qRT-PCR |
| *Homo sapiens* | *HPRT* | TTGGTCAGGCAGTATAATCC | GGGCATATCCTACAACAAAC | qRT-PCR |
| *Homo sapiens* | *KDELR3* | TCTGTACCGGGCACTCTACC | ACTTCTTTCCCTTAAGGACTTTGGT | qRT-PCR |
| *Homo sapiens* | *MTX2* | AGGGGAGATCACTCATGCTAGG | TGACAGCACTGGTCTACATCC | qRT-PCR |
| *Homo sapiens* | *PELP1* | GAGCCCCACAGAGCTATTCC | GGGTCCTGGGTCTGTAACAC | qRT-PCR |
| *Homo sapiens* | *RPS7* | TCGTCTTTATCGCTCAGAGGAG | GCACAGCTGTCAGAGTACGG | qRT-PCR |

| | | | | |
|---|---|---|---|---|
| *Homo sapiens* | *SEC23B* | TACGTGATACAGCGAGGTGC | GCAGGGACTCTTTGAGTGCT | qRT-PCR |
| *Homo sapiens* | *TWIST1* | CGGAGACCTAGATGTCATTGTTT | CCACGCCCTGTTTCTTTGAAT | qRT-PCR |
| *Homo sapiens* | *USP14* | GGCTTCAGCGCAGTATATTA | CAGATGAGGAGTCTGTCTCT | qRT-PCR |
| *Homo sapiens* | *AEBP1* | GCTTACTAATGCGCACGCGA | CGTACAACCACAGCACCAAC | ChIP |
| *Homo sapiens* | *COL4A1* | CAGTGGAACAGAGCTTCGTAAAAC | TATTGAACCAGTGCTGGAAGGAA | ChIP |
| *Homo sapiens* | *COL4A1 (2)* | GCATTGCAAACGCCAGACA | CGTTGGCTGGCTAAATGGGT | ChIP |
| *Homo sapiens* | *FKBP10* | CAACTCCAGGCACCATGTTC | CCTGCACCACCAGTAGCAG | ChIP |
| *Homo sapiens* | *PELP1* | ATCTGAAGTGCTGGCAACCG | GGTCATCTGGAGAACTTCCCTC | ChIP |
| *Homo sapiens* | *SNAI2* | CCGCTTCCCCCTTCCTTTTT | AGCCTCTGGTGTTAATGAGAGC | ChIP |
| *Homo sapiens* | Neg | AGTGCCTGCACCCAAGATTT | TGCAAACCTGCTTAACTCCAAC | ChIP |

**Table S3 Antibodies**

| Antibody | Origin | Dilution | Application | Source | Catalog no. |
|---|---|---|---|---|---|
| TWIST1 | Mouse | 1/200 | WB | Santacruz | sc-81417 |
| TWIST1 | Mouse | 1/200 | IF | Abcam | ab175430 |
| Calnexin | Mouse | 1/3000 | WB | Bd Transduction Laboratories | 4178754 |
| PDHX | mouse | 1/3000 | WB | Santacruz | sc-393644 |
| COL4A1 | Rabbit | 1/1000 (WB/IF), 1/100 (IHC-f) | WB/IF/IHC-f | Genetex | GTX130215 |
| AEBP1 | Rabbit | 1/1000 (WB/IF), 1/50 (IHC-f) | WB/IF/IHC-f | Invitrogen | PA5109366 |
| FKBP65 | Rabbit | 1/1000 | WB/IF | Proteintech | 12172-1-AP |
| PELP1 | Mouse | 1/2000 | WB | Proteintech | 67050-1 |
| Ki67 | Rabbit | 1/200 | IF | Abcam | AB15580 |
| VE-Cadherin | Mouse | 1/300 | IF | BD Biosciences | 555661 |
| CD31 | Rabbit | 1/500 | IF | Abcam | ab182981 |
| anti-Mouse | Goat | 1/500 | IF | Thermofisher | A-11001 |
| anti-Rabbit | Goat | 1/500 | IF | Thermofisher | A-11011 |

| | | | | | |
|---|---|---|---|---|---|
| anti-Mouse (HRP) | Goat | 1/3000 | WB | Agilent/Dako | P0447 |
| anti-Rabbit (HRP) | Goat | 1/3000 | WB | Agilent/Dako | P0448 |
| AF488-CD31 | Rat | 1/50 | FACS | Biolengend | 102514 |
| APC-CD45 | Rat | 1/100 | FACS | Biolengend | 103112 |
| TruStain FcX™ CD16/32 | Rat | 1/50 | FACS | Biolengend | 101320 |
| AF488-ACTA2 | Mouse | 1/200 | IHC-f | Abcam | Ab184675 |
| PELP1 | Rabbit | 1/400 | IHC-f | Invitrogen | PA5-76700 |
| VWF | Rabbit | 1/300 | IHC-f | DAKO | A0082 |
| ACTA2 | Mouse | 1/150 | IHC-p | DAKO | M0851 |
| ACTA2 | Mouse | 1/1000 | IF | DacoCytomation | M0851 |
| MAC3 | Rat | 1/75 | IHC-p | BD-Pharmigen | 550292 |
| Anti-FLAG | Rabbit | 10ug | ChIP | Cell signalling | 14793S |
| IgG | Rabbit | 10ug | ChIP | Diagenode | C15410206 |
